# Predicting bacterial interaction outcomes from monoculture growth and supernatant assays

**DOI:** 10.1101/2023.07.03.547502

**Authors:** Désirée A. Schmitz, Tobias Wechsler, Ingrid Mignot, Rolf Kümmerli

## Abstract

How to derive principles of community dynamics and stability is a central question in microbial ecology. Bottom-up experiments, in which a small number of bacterial species are mixed, have become popular to address it. However, experimental setups are typically limited because co-culture experiments are labor-intensive and species are difficult to distinguish. Here, we use a 4-species bacterial community to show that information from monoculture growth and inhibitory effects induced by secreted compounds can be combined to predict the competitive rank order in the community. Specifically, integrative monoculture growth parameters allow building a preliminary competitive rank order, which is then adjusted using inhibitory effects from supernatant assays. While our procedure worked for two different media, we observed differences in species rank orders between media. We then parameterized computer simulations with our empirical data to show that higher-order species interactions largely follow the dynamics predicted from pairwise interactions with one important exception. The impact of inhibitory compounds was reduced in higher-order communities because their negative effects were spread across multiple target species. Altogether, we formulated three simple rules of how monoculture growth and supernatant assay data can be combined to establish a competitive species rank order in an experimental 4-species community.

## Introduction

Microbial communities are diverse in composition and function (1,2). Top-down approaches based on 16s rRNA and shot-gun sequencing are commonly used to describe community diversity and functional capacities, respectively (3–5). However, much less is known about the microbial interactions occurring in these communities. This is why bottom-up approaches have become popular, whereby a set of defined bacterial species and strains is mixed and their interactions studied under experimental conditions (6–12). The power of simple communities is that they are tractable and allow for experimental manipulation so that the effect of specific factors on community properties can be examined. There is a plethora of bottom-up studies that shed light on various ecological aspects of species interactions. Insights from such studies include, for example, that harsh environmental conditions seem to be more conducive to ecological facilitation (6) and for promoting biodiversity and community stability (7). Furthermore, social interactions also appear to be determinants of community composition and stability, be it through metabolic cross-feeding between species (8,10) or the secretion of competitive and cooperative secondary metabolites (11).

Despite their simplicity, studying experimental communities also comes with several challenges. For example, it is often unclear what the key determinants of interactions are. This is particularly the case for communities for which no a priori knowledge of relevant traits and species characteristics is available. Here, several factors could determine species interactions and competitiveness. The first factor is monoculture growth performance. In a simple environment that is nutrient-rich with low niche diversity, a high growth rate is often taken as a proxy for competitiveness (13,14), and it is expected that fast-growing species have a competitive advantage over slower-growing ones. The second factor involves secreted compounds that could either positively (e.g., amino acids) or negatively (e.g., toxic compounds) affect the growth of other species (7,15–18). The third factor entails interactions that can only arise when species grow together. They can include physical interference through killing systems (e.g., type VI secretion system) as well as niche or metabolic competition whereby species reduce or prevent competitors from accessing certain resources (19–21). A key challenge is to assess the relative importance of these three factors and how they combine to predict species dominance and community dynamics. Further challenges arise when considering that species interactions might differ across environments (6,7) and that pairwise interactions might not necessarily predict interaction dynamics in larger communities (9) due to non-additive effects and new emerging properties.

In our study, we combine experiments and computer simulations on a 4-species bacterial consortium to tackle some of these challenges. Our consortium consists of four opportunistic human pathogens (*Burkholderia cenocepacia* [B], *Cronobacter sakazakii* [C], *Klebsiella michiganensis* [K], *Pseudomonas aeruginosa* [P]), and has been established as a model community in one of our earlier studies (22). With this model consortium, we pursue four main aims. First, we ask whether differences in monoculture growth performance and interactions based on stimulatory or inhibitory molecules secreted by bacteria are good predictors of species interactions in pairwise co-culture experiments. Second, we compare whether the observed interaction patterns are consistent across two different nutritional environments: Lysogeny broth (LB) and Graces’ insect medium (GIM). While LB is a standard laboratory medium, GIM mimics the environment found in Lepidoptera larvae, a common host model system (23–25). Third, we derive tentative rules to predict pairwise interaction outcomes from monoculture growth and supernatant assays and establish rank orders for competitiveness for our 4-species community. Finally, we combine experiments with 3-species and the 4-species communities and agent-based modeling (26,27) (parametrized with our experimental data) to examine whether higher-order community interactions recover the competitive rank orders predicted from pairwise co-culture experiments.

### Materials and methods Bacterial strains

For all experiments, we used the following four bacterial species: *Pseudomonas aeruginosa* PAO1 [P] (28), *Burkholderia cenocepacia* K56-2 [B] (29), *Klebsiella michiganensis* [K], and *Cronobacter sakazakii* (ATCC29004) [C]. We had established this consortium of four gram-negative bacterial species in an earlier study (22), in which we were interested in interactions between pathogens with different virulence levels in the larvae of the greater wax moth *G. mellonella*.

### Monoculture growth assays

All media were purchased from Sigma Aldrich, Switzerland, unless indicated otherwise. All experiments in this study were carried out in lysogeny broth (LB) and Grace’s insect medium (GIM) (Gibco, GIM 1X, supplemented). GIM is a nutrient-rich medium and consists of 19 different amino acids, 10 different vitamins (e.g., biotin, riboflavin, folic acid), 5 inorganic salts (CaCl_2_, MgCl_2_, MgSO_4_, KCl, NaH_2_PO_4_-H_2_O), and several carbon sources (e.g., fructose, glucose, sucrose). Before experiments, each species was taken from a 25% glycerol stock stored at -80° C and grown overnight in 5 mL LB at 37° C and 170 rpm with aeration (Infors HT, Multitron Standard Shaker) until stationary phase. The bacterial cultures were then washed twice with 0.8% NaCl, centrifuging for 5 min at 7,500 rcf (Eppendorf, tabletop centrifuge MiniSpin plus with rotor F-45-12-11). Subsequently, we adjusted the cultures to an optical density at 600 nm (OD_600_) = 1. Then, each well of a 96-well cell culture plate (Eppendorf, non-treated, flat bottom) was filled with 190 μL medium (LB or GIM) and 10 μL individually diluted cell suspensions to reach a starting number of 10^5^ CFU/mL (i.e., 2×10^4^ CFU/well). Plate layouts (i.e., positions of species and replicates on plates) were randomized between experiments. To minimize evaporation, the outer moat space and inter-well spaces of the microplate were filled with 13 mL sterile water. Plates were incubated at 37° C in a microplate reader (Tecan, Infinite MPlex) and OD_600_ was measured every 15 min for 36 h with 60 sec shaking before each reading. We used the Gompertz function to fit growth curves to the obtained OD_600_ trajectories and to extract three different growth parameters: the maximum growth rate (μ_max_), area under the curve (AUC), and inverse of the time to mid-exponential phase (1/T_mid_). For the latter, we used the inverse of T_mid_ to ensure that low and high values stand for low and high growth, respectively. The parameter 1/T_mid_ considers the lag phase λ, μ_max_, and yield A and was calculated as:

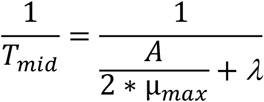

### Supernatant assays

Cells were prepared as for the monoculture growth assays. Washed cell cultures were subsequently inoculated in 10 mL medium within 50 mL Falcon tubes (inoculum size: 10^5^ CFU/mL). We had 10 replicates (tubes) per species. Tubes were incubated at 37° C shaken at 170 rpm with aeration (Infors HT, Multitron Standard Shaker). We harvested supernatants when species had reached the stationary phase, which was after 16 hours for C and P, and after 40 h for B and K (Fig. 1). Supernatants from all 10 replicates of a species were pooled and mixed before filter sterilizing them with a 0.2 μm filter (GE Healthcare Life Sciences, Whatman filter) and aliquoting and storing them in 2 mL Eppendorf tubes at -20° C.

**Figure 1.**
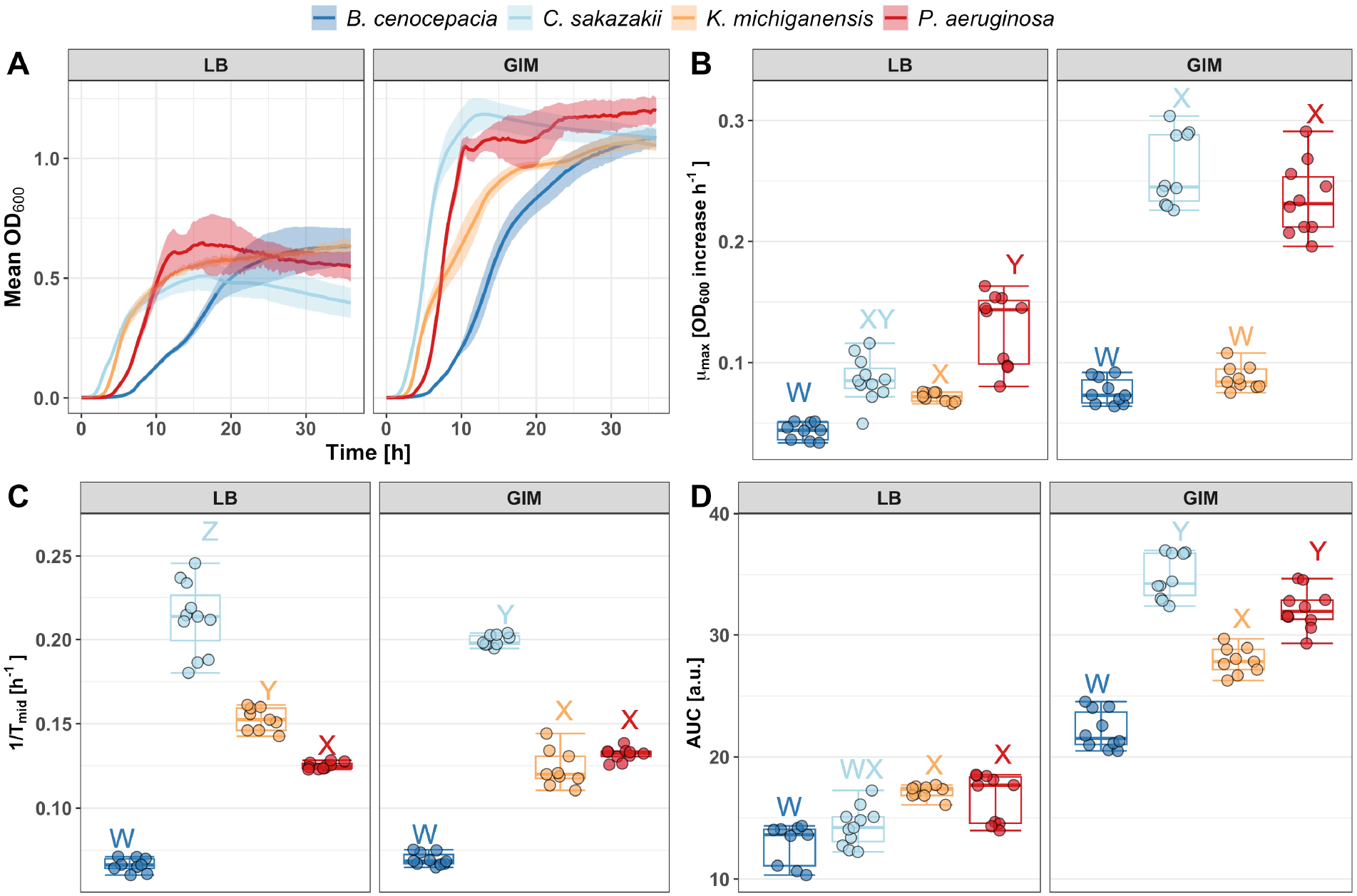
Growth differences of the four bacterial species in two different media: LB medium and Grace’s insect medium [GIM]. (A) Growth curves over 36 h in shaken, liquid cultures. Shaded areas depict the standard deviation. Boxplots show the (B) maximum growth rate μ_max_ as OD_600_ increase per hour (C) the inverse of the time to mid-exponential phase [1/T_mid_], and the (D) integral [area under the curve, AUC]. All growth parameters are derived from the growth curves in (A) using the Gompertz curve fit. Different letters above boxplots indicate significant growth differences (alpha = 0.05) between bacterial species using linear mixed models with experimental block as random factor. Boxplots show the median (line within the box) with the first and third quartiles. The whiskers cover 1.5x of the interquartile range or extend from the lowest to the highest value if all values fall within the 1.5x interquartile range. Data are from 3 independent experiments, each featuring 3-4 replicates per condition, resulting in a total of 9-10 replicates per condition.

For the actual supernatant assays, we prepared cells and distributed them on microplates as described for the monoculture growth assay. The cells were then exposed to three different media: (a) 30% supernatant from another species + 70% fresh medium (LB or GIM), (b) 30% NaCl (0.8% solution) + 70% fresh medium, and (c) 100% fresh medium. Plates were incubated at 37° C in a microplate reader (Tecan, Infinite MPlex) and OD_600_ was measured every 15 min for 36 h with 60 sec shaking before each reading. We fitted Gompertz functions to the growth trajectories and extracted μ_max_, AUC, and 1/T_mid_. We then expressed the growth in supernatant relative to the two control treatments, (a)/(b) and (a)/(c). We define growth inhibition if (a)/(b) < 1, and growth stimulation if (a)/(c) > 1.

To obtain a comparable species ranking across assays, we calculated an integrative supernatant score. We summed up all the relative growth values (a)/(b) that the supernatant of one species has on the other three species. The four values (one per species) were then centered around 0 and scaled between -1 and +1. Negative and positive values stand for stimulatory and inhibitory supernatant effects, respectively, and are indicative of a species’ competitiveness.

### Co-culture experiments

Overnight cultures were washed and OD_600_ was adjusted as described for the monoculture growth assays. Next, bacterial species were diluted individually to reach similar cell numbers. To obtain 10^5^ CFU/mL per well for pairwise co-cultures (i.e., 5×10^4^ CFU/mL per species), 25 μL from each of the two species’ dilutions was added in each well of a 24-well tissue culture plate (Corning, flat bottom with low evaporation lid) filled with 950 μL medium. For 3-species and 4-species communities, the same total number of 10^5^ CFU/mL was added per well, with equal amounts of each species. Well-plates were incubated at 37° C and 170 rpm for 24 h (Infors HT, Multitron Standard Shaker).

After competition, OD_600_ was measured to calculate the appropriate dilution for plating. Calculations were based on previously established calibration curves, relating OD to CFU for each of the four species and averaging the expected CFU between the two species in a mix. Cultures were diluted with 0.8% NaCl, and each replicate was plated in duplicate on LB-agar plates (1.5% agar). We incubated all plates overnight at 37° C and manually counted CFU on the next day. All four species could be distinguished from each other based on their different colony morphologies on plates (22). Since C colonies turn yellow after a day at room temperature, we left them on the bench for an additional day. Plates containing B were incubated at 37° C for two days in total because of its slow growth. If one species was much more frequent than the other one, we used selective plating (if available) to ensure that we could assess the abundance of the low-frequency species. For P, we used Pseudomonas isolation agar.

For B, we used LB plus gentamicin (30 μg/mL) due to its inherent resistance to this antibiotic. No selective media was available for C, meaning that C could not be counted in co-culture with P below a given detection limit.

For pairwise co-cultures, the relative fitness values of each focal species were calculated based on Wrightian fitness from Ross-Gillespie et al. (2007) as follows:

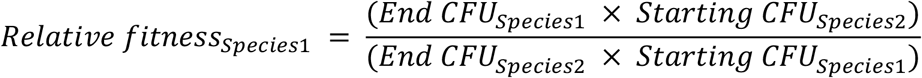

We log-transformed the relative fitness data to be able to better compare values across species combinations. Relative fitness values > 0 and < 0 indicate that the focal species wins or loses in competition with the co-cultured species, respectively. For the 3-species and 4-species communities, the above formula cannot be applied and we thus directly compared CFUs across species after competition.

For pairwise co-cultures, we calculated an integrative score to obtain a measure of competitiveness for each species relative to all other species. Given that each relative fitness value can be expressed from the perspective of the winner or the loser (resulting in reciprocal values), we followed a stepwise process to avoid double counting of fitness values. We started with the weakest species and summed up all its relative fitness effects in competition with the three other species. Then we moved to the second (and third) weakest species and repeated the procedure leaving out any fitness effects that were already accounted for before. For the final two species, we only considered their effect on each other. With this approach, opposing fitness effects can balance each other out. For example, a species of interest might have a positive effect on one species but a negative effect on another species, such that it will take up a relatively neutral position in the ranking.

### Agent-based model

We performed agent-based simulations, using our previously developed platform (26,27). Microbial simulations take place on a two-dimensional toroidal surface with connected edges without boundaries. The surface of the torus is 10,000 μm^2^ (100 × 100 μm). Bacteria are modeled as discs with an initial radius of 0.5 μm. Bacteria grow through an increase of their radius, according to a growth function, and divide when reaching the threshold radius of 1 μm. In our simulations, we assumed that resources are not limited. Growth differences emerge solely based on the differences in growth rate and interactions between the species. Interactions are modeled via secreted molecules. Molecules can be taken up when they physically overlap with a cell and can either be growth-stimulating or -inhibitory. Here, we only modeled inhibitory molecules (represented by a toxin, because all major supernatant effects were negative in GIM, see Fig. 2). Accordingly, the growth of each cell is determined by the function

**Figure 2.**
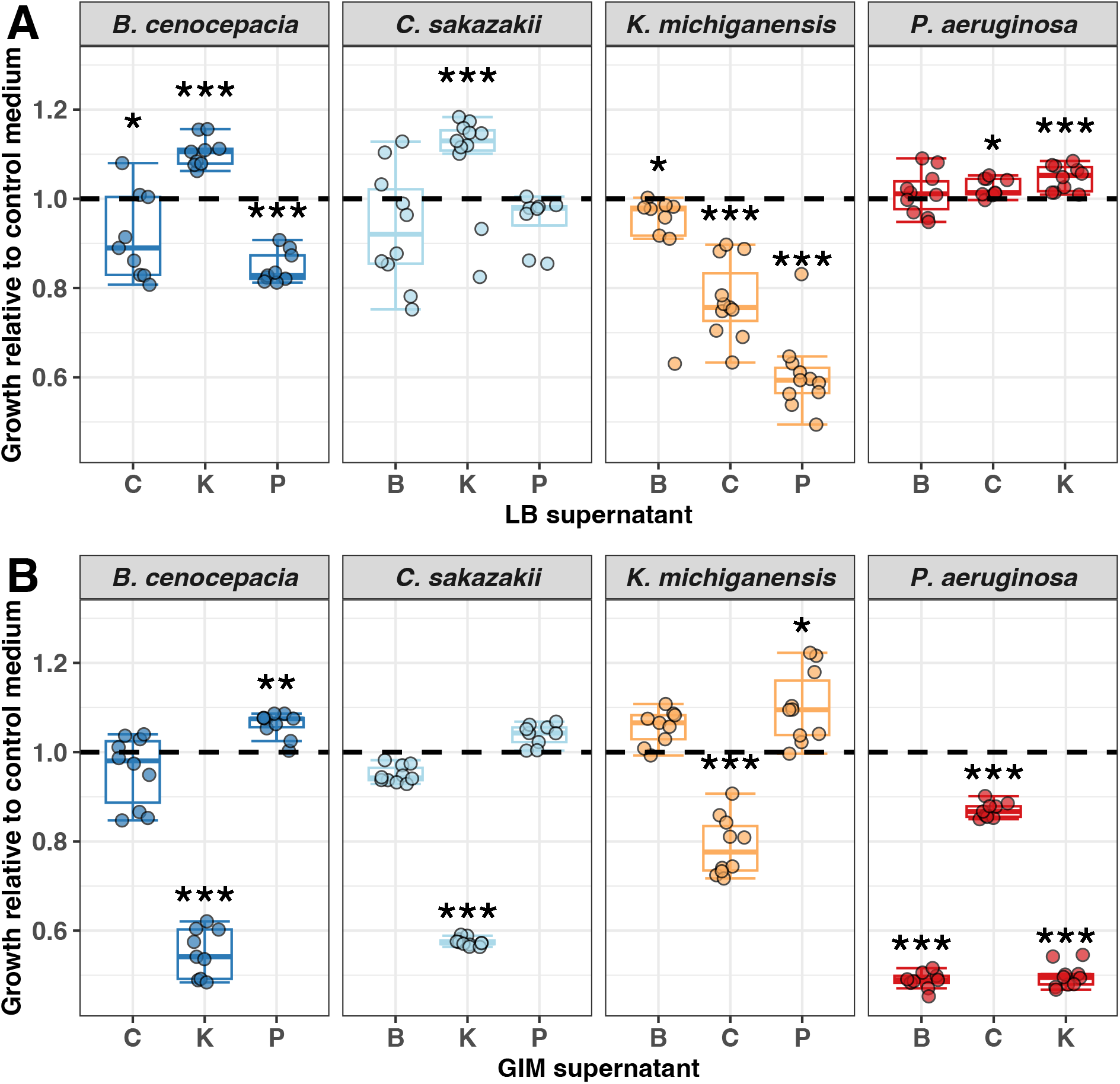
Boxplots show the relative growth of each species in the conditioned medium (30% supernatant + 70% fresh medium) of the other species both in (A) LB and (B) GIM medium, compared to a control treatment, depicted by the black dashed line (30% NaCl solution [0.8%] + 70% fresh medium) measured across 36 h of growth. For this plot, we used 1/T_mid_ for growth comparisons (see Fig. S3 for absolute readouts). We repeated the same analysis for μ_max_ (Fig. S4, S5) and AUC (Fig. S6, S7). Relative growth was calculated by dividing the absolute 1/T_mid,_ (estimates from curve fits) in the supernatant treatments by 1/T_mid_ in the control treatment. Asterisks depict significant differences (alpha = 0.05) of a species’ growth in the particular supernatant compared to its growth in the control medium using a linear mixed model with experimental block as random factor. Boxplots depict the median (line within the box) with the first and third quartiles. The whiskers cover 1.5x of the interquartile range or extend from the lowest to the highest value if all values fall within the 1.5x interquartile range. Data are from 3 independent experiments, each featuring 3-4 replicates per condition, resulting in a total of 9-10 replicates per condition. The underlying growth curves can be found in the supplementary information (Fig. S8).

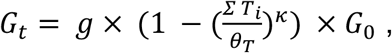

where *G*_*t*_ denotes the increase in radius per time step, *G*_0_ denotes the default radius increase per time step (0.5/1, 200 μm), *g* the species-specific growth rate, and *ΣT* the number of accumulated toxins. Toxins decrease growth and lead to cell death when they accumulate beyond the threshold value *θ*_*T*_. The decrease in growth is further controlled by the latency parameter *κ* = 2, which leads to an exponentially increasing (negative) effect on growth, depending on the number of accumulated toxins. If a cell divides before the threshold *θ*_*T*_ is reached, the accumulated toxins are shared equally among the two daughter cells.

Simulations started with 32 cells per species, randomly placed on the landscape. Bacteria can disperse on the surface, according to a specific cell diffusion coefficient *D*. Bacteria start to grow and divide until a cell number of 500 is reached. A chemostat mechanism is then activated to keep cell numbers constant at around 500 by randomly removing cells. The chemostat mechanism allows observing strain dynamics over extended timespans and prevents surface overgrowth. We ran simulations for 30,000-time steps with 20 replicates per parameter and species combination.

To implement differences in growth rates between the four species, we used scaled differences in 1/T_mid_ from monoculture growth in GIM. The specific growth rate of each species is

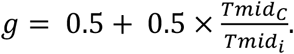

where Tmid_C_ and Tmid_i_ are the growth parameters of the best-growing species C and the other species, respectively. Accordingly, the species-specific growth rates were C = 1, P = 0.84, K = 0.81, and B = 0.68.

We allowed strains to produce toxins to capture the inhibitory effects observed in the supernatant assays in GIM. In practice, the inhibitory effects could be caused by other factors than toxins, such as secreted molecules that deplete a specific nutrient (e.g. siderophores reducing iron availability) or induce a change in pH (e.g., P supernatant slightly increased pH from 6.2 to 6.0 in GIM, Table S1). Thus, the toxin implemented here is representative of any type of molecule that negatively affects the growth of another species. We only considered effects affecting growth by at least 20%. This was the case for K inhibiting all other species, and B inhibiting P. To calibrate toxin potency *θ*_*T*_ (defined by the number of molecules required to kill a cell), we simulated pairwise competitions between all four species across a range of values (*θ*_*T*_: 750 to 4,500, in steps of 250). The aim was to define the minimal toxin potency required to recover the competitive outcomes observed in the co-culture experiments. We found this to occur with the following values: K-toxin against C, *θ*_*T*_=1,750; K-toxin against B and P, *θ*_0_=2,500; B-toxin against P, *θ*_*T*_=2,500 (Fig. S1). Note that low values stand for high toxic potency because fewer toxin molecules are needed to kill a cell. These simulations were conducted with high cell and molecule diffusion (*D* = 5 μm^2^s^−1^ and ∂ = 10 μm^2^s^−1^ respectively) to match experimental shaken culture conditions.

Following calibration, we modeled higher-order competitions between all 4-species and all possible 3-species combinations. For these simulations, we considered environments with different diffusion conditions (low: D = 0.0 μm^2^s^−1^, ∂ = 0.1 μm^2^s^−1^, high: *D* = 5.0 μm^2^s^−1^, ∂ = 10.0 μm^2^s^−1^). For each replicate, we extracted species frequency at each time point, calculated the mean species frequency over time, and recorded the per capita toxin uptake per species.

### Statistical analyses

All statistical analyses were conducted with R (version 4.1.1) and RStudio (version 2021.09.0+351). We built linear mixed models with experimental date as a random variable (i.e., experiments were repeated in independent blocks on different days), for both the monoculture growth and supernatant assays. We fitted the growth term as response variable and species (monoculture) or medium (supernatant) as explanatory variable. We used the function “transformTukey” (package “rcompanion” (30)) to find the best transformation of the response variable to meet normally distributed residuals. To adjust p-values in multiple pairwise comparisons, we used the R function “pairs” from the “emmeans” package (31) based on the Tukey method for monoculture growth data, and the FDR method for supernatant data. To test whether the relative fitness in pairwise competitions is different from the expected log(1) = 0 value (assuming that both species perform equally well), we used one-sample two-sided Wilcoxon rank tests, since our relative fitness data were not normally distributed. We used Wilcoxon signed rank tests for pairwise comparisons in 3- and 4-species communities, adjusting p-values with the FDR method. All details of the statistical analyses can be found in a separate statistics file.

## Results

### Monoculture growth parameters as predictors of competitiveness

In a first experiment, we assessed the growth performance of the four pathogen species in monoculture in our two media (LB and GIM) over 36 hours. From the obtained growth curves, we estimated the maximum growth rate (μ_max_), the inverse of the time to mid-exponential phase (1/T_mid_), and the growth integral (area under the curve, AUC).

All species grew better in GIM than in LB, and growth trajectories differed between the four species (Fig. 1A). When focusing on μ_max_, we observed that C and P grew better than B and K, with the differences being larger in GIM than in LB (Fig. 1B). Next, we looked at 1/T_mid_ and found that this growth parameter returns C and B as the best and worst grower, respectively, but downgrades the performance of P relative to μ_max_ because of the relatively long lag phase of P. The AUC dampened the differences between species (Fig. 1D). This is expected as the integral gives more weight to yield, which is quite similar for all four species.

Our analysis reveals that the competitiveness of a species derived from monoculture readouts depends on the growth parameter examined. This raises an additional challenge, namely which growth parameter to consider. To address this challenge, we conducted correlation analyses (Fig. S2) and found that all three growth parameters correlate well with each other in GIM (Pearson correlation coefficients, r = 0.81 to 0.91). However, this is not the case in LB, where correlations are poor or non-significant (r = 0.33 to 0.56). The main reason for these mismatches are P with its high μ_max_ but low 1/T_mid_, and C with an intermediate μ_max_ but high 1/T_mid_ due to a short lag phase (Fig. 1). Since we cannot yet infer which parameter predicts species competitiveness best, we estimated all three growth parameters for both media and show the species rank order for all of them (Fig. 4).

### Effect of secreted compounds as predictors of competitiveness

We used supernatant assays to assess whether secreted compounds can predict competitiveness between interacting species. Supernatants contain molecules (e.g., toxins, siderophores, biosurfactants, amino acids) secreted by the producing species that can negatively or positively impact the fitness of the receiving species (7,15–18).

To test for such effects, we exposed each of our species to the supernatant (30% supernatant + 70% fresh medium) of all other species and measured the growth effects relative to a control condition (30% NaCl solution [0.8%] + 70% fresh medium). In Figure 2, we present the results with the growth parameter 1/T_mid_, while the results for μ_max_ and AUC are shown in Fig. S3 and Fig. S4, respectively.

Similar to the monoculture parameters, we found differences in the supernatant effects between the two media. While there were 5 negative growth effects out of 12 in LB (Fig. 2A) and 6 negative effects out of 12 in GIM (Fig. 2B), 9 out of the 12 interactions changed direction from neutral/positive to negative (or vice versa) between media. For example, the supernatant of K was highly inhibitory for B, C, and P in GIM but not in LB. Overall, we found that supernatants from one species can have strong inhibitory effects on the growth of another species. We also observed positive supernatant effects. Such stimulatory effects could be caused by several different factors, including nutrient leftovers in the supernatant, or nutrient release and the secretion of beneficial compounds such as enzymes by the species producing the supernatant.

To obtain an integrative score of how much our four species influence each other’s growth indirectly via their supernatant, we summed up all the effects one species has on the others, separately for each of the three growth parameters (i.e., from Fig. 2, S3, and S4). This score yielded species rank orders that differ between growth parameters and media (Fig. 4).

### Competitiveness in co-culture assays

Next, we carried out co-culture experiments in all pairwise species combinations in both media and determined the relative fitness of the competing species after 24 h (Fig. 3). Unlike the previous competition metrics, we found the results to be much more consistent between LB (Fig. 3A) and GIM (Fig. 3B). The competitive outcomes only changed in 2 out of the 6 comparisons and were pronounced in only one case, in which C lost against P in LB but was the clear winner in GIM. An example of consistency is B, which was outcompeted by all competitors in both media. Furthermore, we found that species coexisted in most cases (Fig. S9, showing the absolute readouts underlying the relative fitness measures in Fig. 3). The only exception occurred in LB, where P seemed to displace B and C in most replicates. We again used an integrative score to obtain a measure of competitiveness for each species (Fig. 4) (see Methods). Important to note is that coexistence was assessed after 24 h. By this time, the winning species can be identified, but the measured frequencies might not represent a stable equilibrium of coexistence.

**Figure 3.**
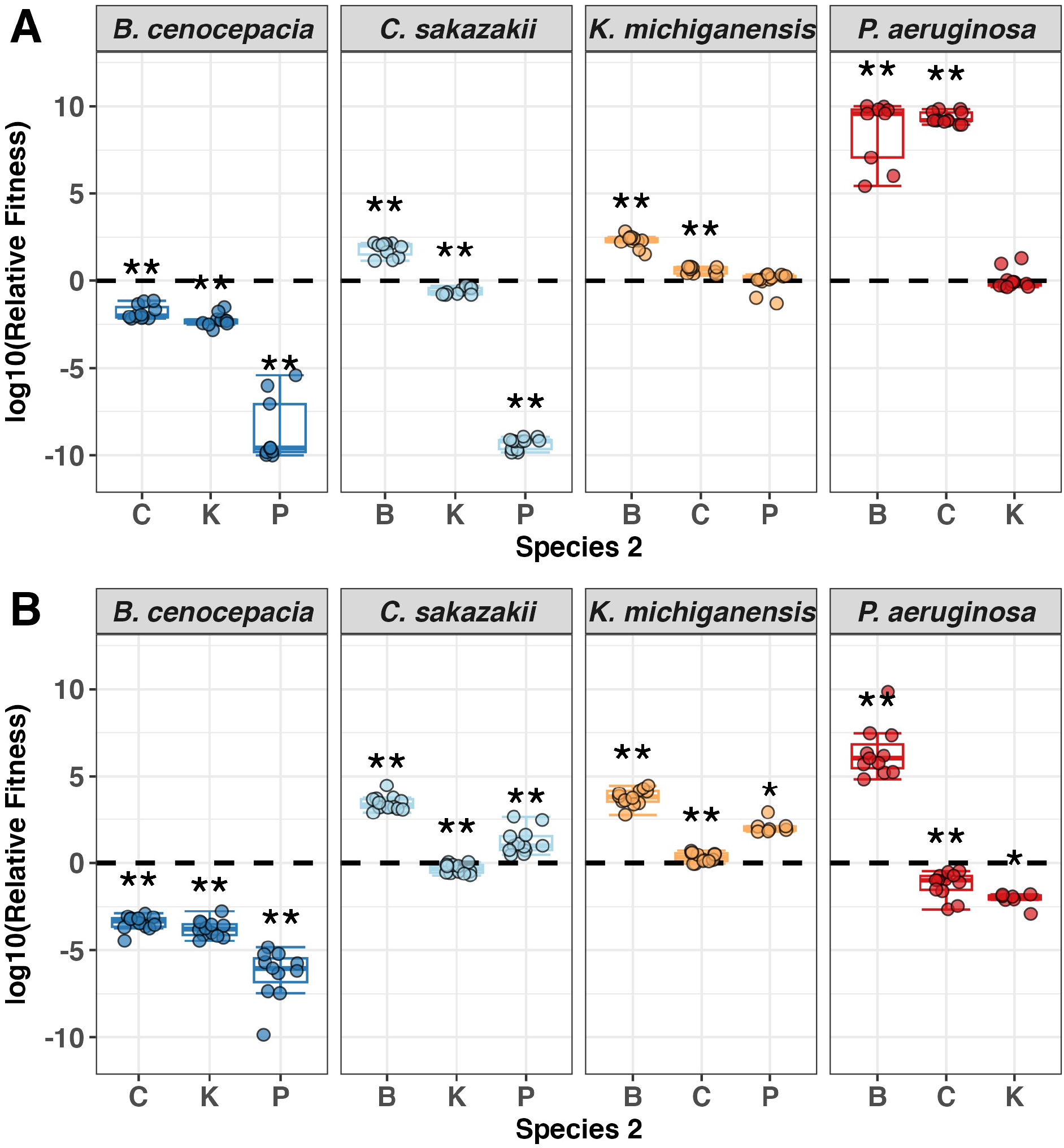
Boxplots show the fitness of the species indicated in the header relative to all other species in co-culture assays in (A) LB and (B) GIM medium. Asterisks depict significant differences (alpha = 0.05) of a species’ fitness relative to that of a competitor against the null hypothesis that none of the two species has an advantage (relative fitness = 0, black dashed line) using one-sample two-sided Wilcoxon rank tests. At relative fitness = 0, two species coexist at equal frequency. Boxplots show the median (line within the box) with the first and third quartiles. The whiskers cover 1.5x of the interquartile range or extend from the lowest to the highest value if all values fall within the 1.5x interquartile range. Data are from 6 individual experiments with 2 replicates per condition, resulting in a total of 7-12 replicates per condition (note in a few cases sample size was <12 because obtaining countable colonies for both species in co-culture was very difficult). The corresponding figure with the absolute readouts can be found in the supplementary information (Fig. S9).

**Figure 4.**
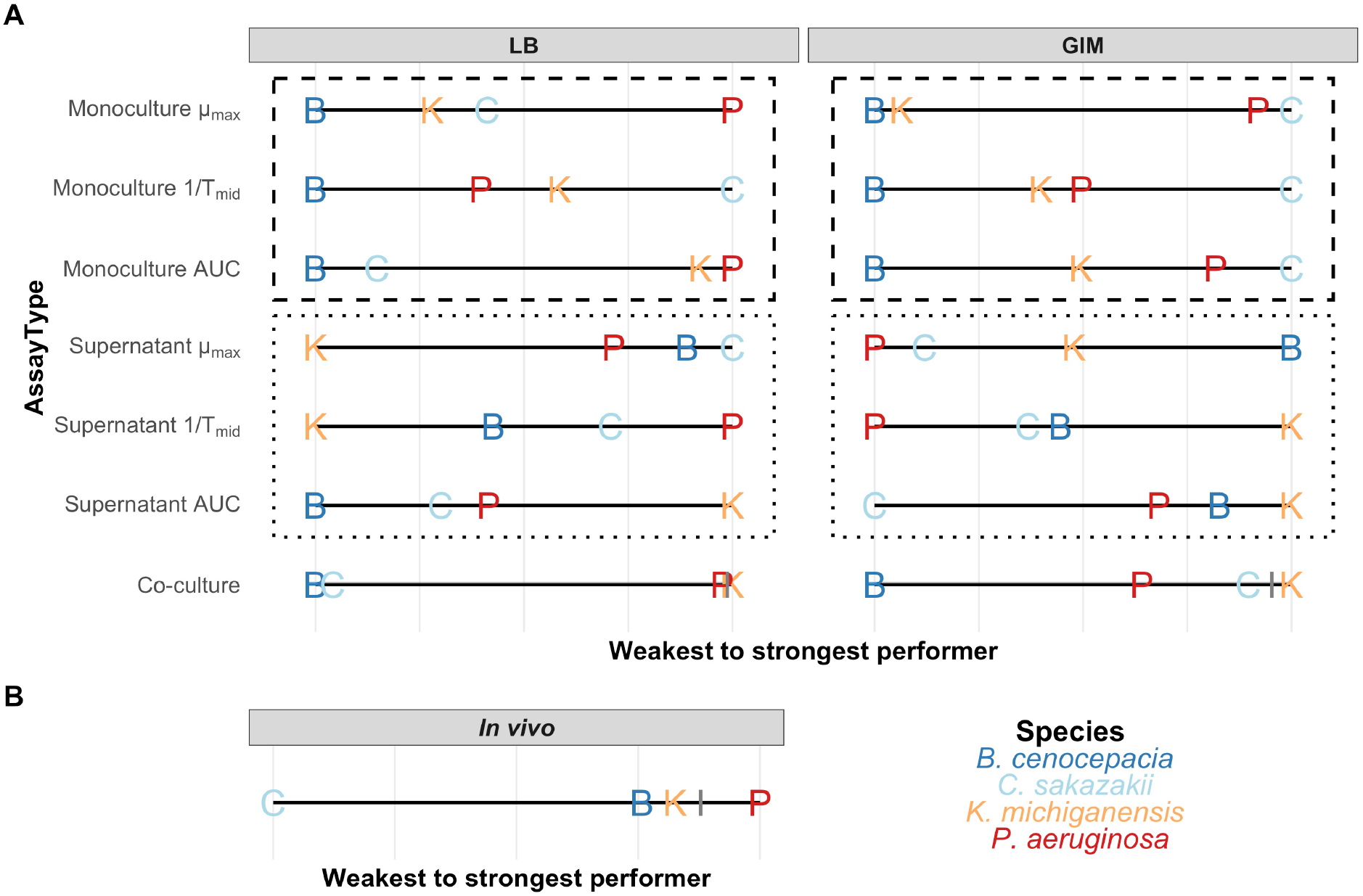
(A) Species rank orders of competitive strength inferred from monoculture growth, the ability of a species to affect the growth of competitors in conditioned medium (supernatant), and co-culture experiments. All *in vitro* experiments were conducted in two different media: LB and GIM. For monoculture growth and supernatant assays, rank orders are shown for maximum growth rate (μ_max_), the inverse of the time to mid-exponential phase (1/T_mid_) and the growth integral (area under the curve, AUC). For the supernatant assays, we summed up all growth effects a species has on the other species and scaled them across species. To scale relative fitness values in co-culture assays, we implemented a stepwise process to avoid double-counting of the reciprocal fitness value. We started with the weakest species and summed up all its relative fitness values. Then we moved to the second (third) weakest species and repeated the procedure leaving out any fitness effects that were already accounted for before. For all experiments, we scaled the values from weakest (left) to strongest (right) performer. A line demarks the transition from neutral/positive to negative effects on the other species. (B) Species rank order from *in vivo* competition experiments in the larvae of *G. mellonella* 12 hours post-infection from a previous study (22). The latter includes all raw data, while Fig. S10 depicts the relative fitness values underlying this figure. The calculation of the species rank order was done as for the co-culture assays. Both co-culture and *in vivo* rank order calculations are based on log10(CFU/mL) values. For more information regarding all calculations, please refer to the methods and Table S4 in the statistical analysis file.

### Deriving tentative rules to predict species competitiveness based on monoculture growth and supernatant effects

In this section, we integrate the competitiveness rankings of the four species across all our experiments and ask which metric or combination of metrics is most predictive of the observed outcome in co-culture experiments (Fig. 4).

Starting with GIM, we find that all monoculture growth parameters positively correlate with one another and return the same species ranking: B < K < P < C. This species ranking matches the outcome of the co-culture experiments well for three species (B < P < C), but not for K. In monoculture, K shows intermediate growth performance but is the most competitive species in co-culture experiments. To explain this mismatch, we consider the supernatant assay results (Fig. 2) showing that K strongly inhibits all other species (based on 1/T_mid_ and AUC). Thus, we can derive two tentative rules to predict competitive outcomes in GIM. Rule 1: take the species rank order based on monoculture growth (B < K < P < C). Rule 2: consider the supernatant effects and adjust the rank order by moving species that strongly inhibit others further up the ranking (B < P < C < K). Interestingly, we find that B (the weakest performer in monoculture) strongly inhibits P in the supernatant assay (Fig. 4), but this inhibition did not lead to a shift in the species ranking in the co-culture experiments. We can thus derive a third tentative rule. Rule 3: inhibitory supernatant effects exerted by slow-growing species can be ignored probably because their slow growth limits sufficient toxic compound production.

For LB, we realize that the species rankings are different for all three monoculture growth parameters and the corresponding supernatant effects. Nonetheless, we applied our three tentative rules to LB and found a good fit when using the AUC growth parameter. Rule 1: take the monoculture species rank order. For LB (AUC), B < C < K < P. Rule 2: consider the supernatant effect and move species that inhibit others further up the ranking (for LB (AUC): first rank: K inhibits B and C / second rank: P inhibits C and K). In this case, the supernatant effects do not induce a major change in the ranking, but a strong clustering into two inferior and two superior species: B ≈ C < P ≈ K. Rule 3: ignore inhibitory supernatant effects of species with poor monoculture growth. For LB (AUC), this rule applies to C, which grows poorly yet inhibits B. Applying these rules leads to a species ranking for LB co-culture competitions that matches our experimental observations. Based on the LB data, we may further derive that more integrative growth measures, such as 1/T_mid_ and AUC might be more informative for competitiveness prediction than growth parameters capturing only a single feature of growth dynamics (μ_max_).

### Can within-host species interactions be predicted from *in vitro* experiments?

Competitive metrics obtained from *in vitro* experiments are often taken as proxies to predict interactions in polymicrobial infections. For example, *P. aeruginosa* typically outcompetes *Staphylococcus aureus in vitro* (32–34), a finding used to explain the prevalence of *P. aeruginosa* in co-infections. Here, we ask whether a direct translation of *in vitro* results to the host context is warranted. This is possible because we used the same pathogens here and in our co-infection study with *G. mellonella* insect larvae (22) (see Fig. S5 for methods and recapitulation of the in-host competition data).

When focusing on GIM medium, which mimics the nutritional conditions of the insect environment, we find a moderate match between the rank orders observed in the *in vivo* and *in vitro* (Fig. 4). While K and P were among the stronger competitors in both environments, the weakest agreements were observed for C and B. In GIM, C performed well whereas it was the poorest competitor in the *G. mellonella* host. Conversely, B was the weakest competitor in GIM, yet was competitive against both C and K in the host. This comparison suggests that host-pathogen interactions but also the host environment itself (e.g. spatial structure, biotic and abiotic factors) matter, highlighting that *in vitro* assays might not be adequate predictors of *in vivo* dynamics.

### Simulating higher-order interactions

Next, we parameterized an agent-based model with data obtained from monoculture growth and supernatant assays to test whether simulated 3- and 4-species community interactions yield the competitive rank order inferred from our pairwise co-culture experiments. For this, we used 1/T_mid_ data in GIM and implemented toxins to model susceptibility to secreted molecules (see methods for details).

We simulated the 4-species community for 30,000-time steps in a high diffusion environment in which bacterial agents and toxins move readily (mimicking shaken liquid conditions), and a low diffusion environment in which bacterial agents do not move and toxins diffuse slowly (mimicking surface-attached growth). Our simulations revealed competitive rank orders for the low diffusion (B < P = K < C) and high diffusion (B < P < K < C) environments (Fig. 5A) that differ from the experimentally predicted rank order (B < P < C < K) (Fig. 4A). Notably, C and K swapped places, indicating that the toxic compound of K that was so potent to suppress C in experimental pairwise competitions showed reduced efficacy in the simulated 4-species community.

**Figure 5.**
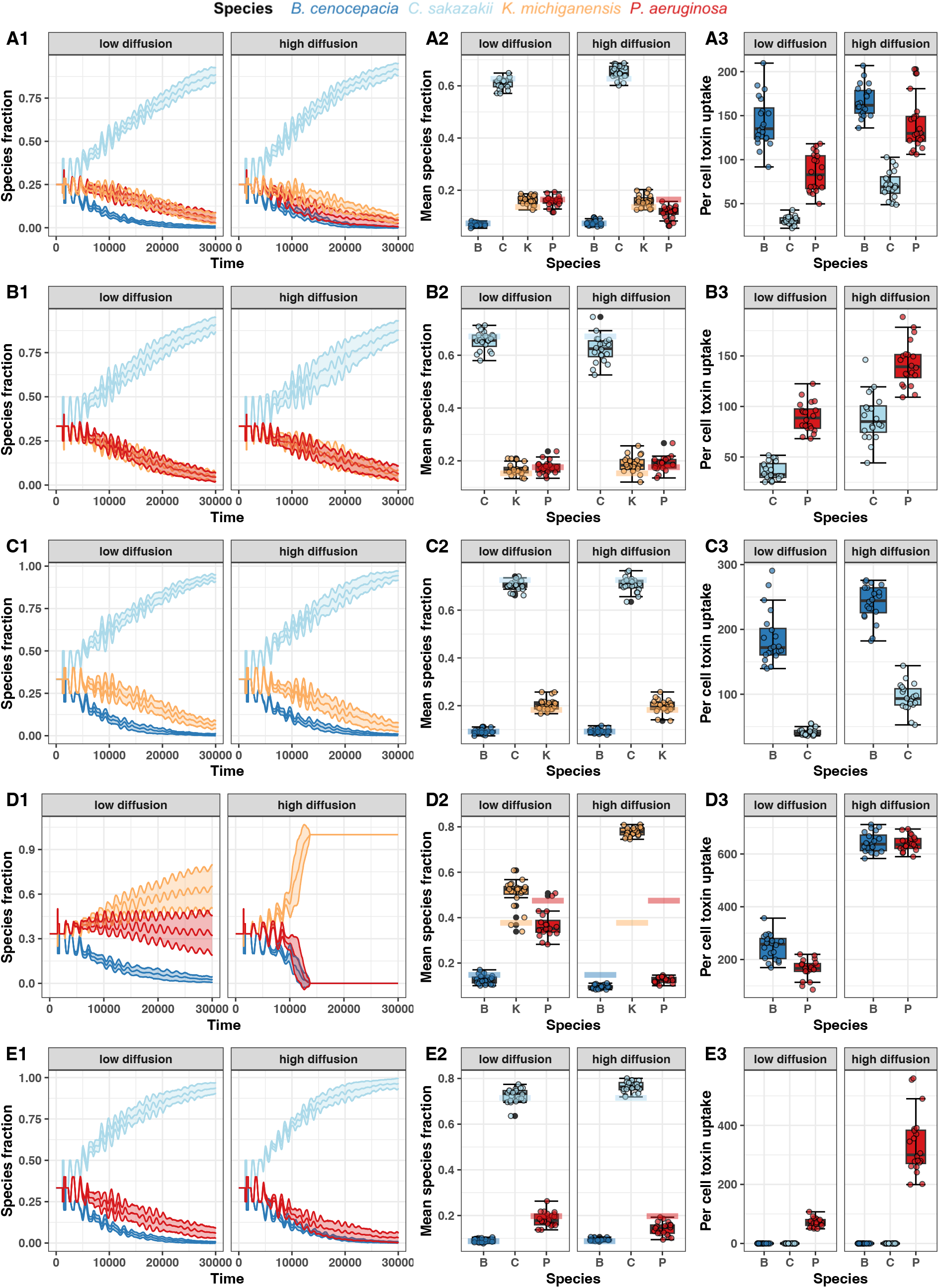
Bacterial community dynamics simulated with an agent-based model and parameterised with experimental growth and inhibition data. Panels show the (1) species fractions over time, (2) mean species fraction over time, (3) per cell toxin uptake for (A) the 4-species community and (B-E) all combinations of 3-species communities. All simulations were carried out under low diffusion (cell diffusion D = 0.0 μm^2^ s^−1^, toxin diffusion ∂ = 0.1 μm^2^ s^−1^, a structured environment) and high diffusion (cell diffusion D = 5.0 μm^2^ s^−1^, toxin diffusion ∂ = 10.0 μm^2^ s^−1^, an unstructured environment). In (1), lines show the mean and the standard deviation across 20 independent simulations. Boxplots show the median values (line within the box) across the 20 simulations with the first and third quartiles. The whiskers cover 1.5x of the interquartile range or extend from the lowest to the highest value if all values fall within the 1.5x interquartile range. In (2), the shaded boxes depict the simulated mean species fraction in the absence of toxins.

How can this non-additive effect be explained? When looking at the toxin uptake rate, we observed that the slow-growing species B and P had the highest per capita toxin uptake rate (Fig. 5A). This suggests the presence of a toxin absorption effect, whereby B and P detoxify the environment for the fast-growing C. A similar case of such an effect has been previously described (35). In our case, the toxin burden is distributed across all susceptible species, yet the fast-growing C benefits the most because it can dilute the cellular toxin concentration below the killing threshold due to its fast replication. In contrast, the slow-growing species reach the toxin threshold and die.

When exploring the 3-species communities (Fig. 5B-E), we found that the presence of a single slow-growing species (either P or B) is enough to create the toxin absorption effect (Fig. 5B+C). We further observed that the toxin absorption effect can be circumvented by higher toxicity, but only when K’s toxin was extremely potent and only under high diffusion conditions (Fig. S6). Importantly, toxin absorption had no effect in the B+K+P community, where K is the fastest-growing species (Fig. 5D). In this community, spatial structure had an important effect by reducing toxin uptake rates and allowing species to coexist. Altogether, our simulations suggest that the toxin absorption effect and spatial structure attenuate toxin efficacy in multispecies communities.

### Experiments with higher-order communities confirm the competitive rank order predicted by simulations

To validate the simulation results, we conducted co-culture experiments with 4-species and 3-species communities in GIM and enumerated CFUs after 24h of competition (Fig. 6). Our experiments recovered the order of competitiveness predicted by the simulations for all five higher-order communities (compare Fig. 5 to Fig. 6). B is weakest, followed by P and K, and C at the top. However, the magnitude of the experimental differences was smaller than the ones observed in the simulations. For example, while K is in second place behind C in both experiments and simulations, K is much closer to C in the experiments than in the simulations. One obvious explanation for this difference is that simulations run much longer than the experiments, magnifying the difference between the two species. Alternatively, the toxin absorption effect may be weaker in experiments as compared to simulations. Altogether, our results demonstrate the usefulness of modeling (36) and its power to predict bacterial community dynamics.

**Figure 6.**
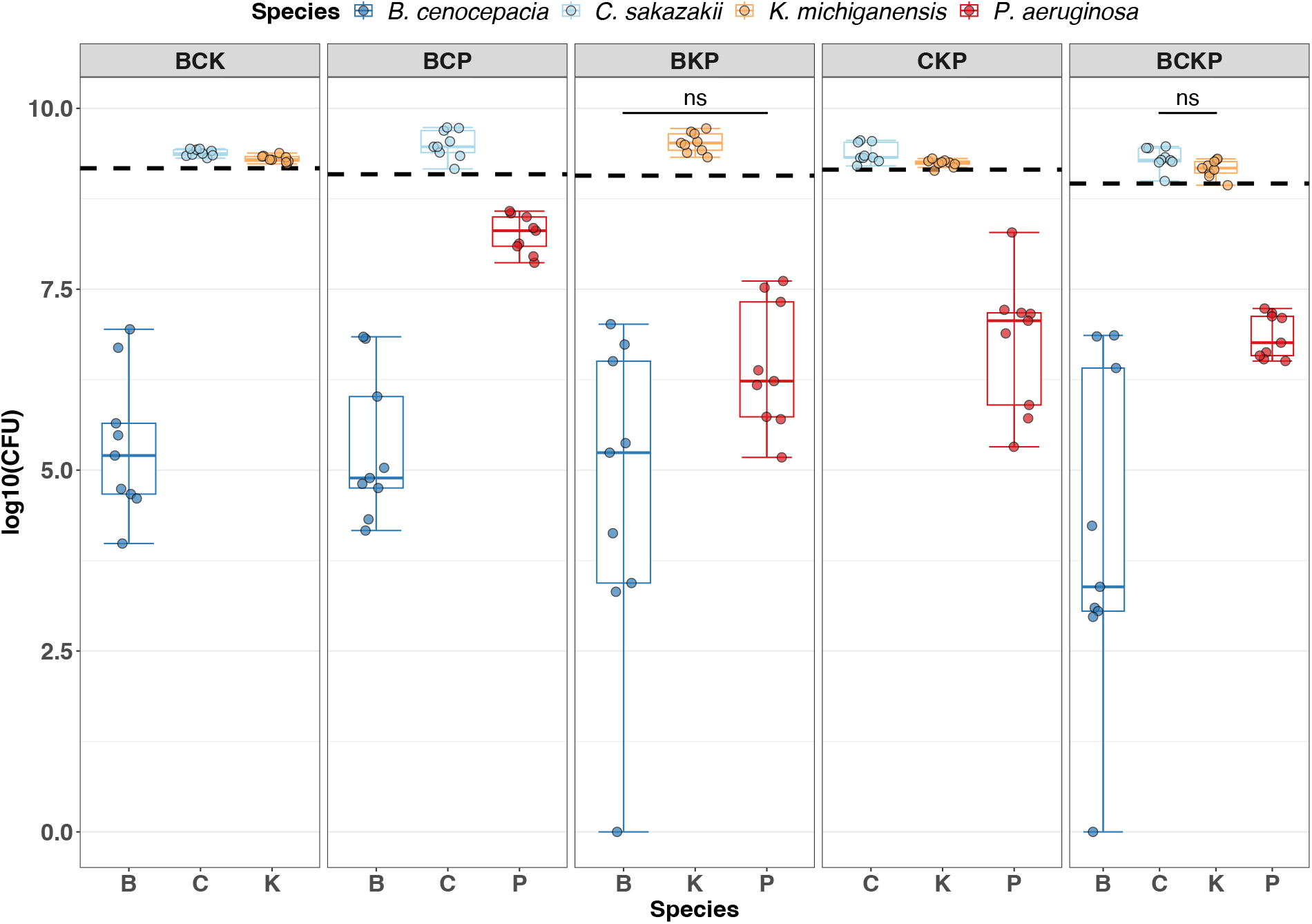
Boxplots show the number of cells of each species in the 4-species and all 3-species communities of our bacterial consortium. Different letters above boxplots indicate significant differences (alpha = 0.05) between bacterial species using Wilcoxon paired signed rank tests for multiple pairwise comparisons with p-value adjustments using the FDR method. Boxplots show the median (line within the box) with the first and third quartiles. The whiskers cover 1.5x of the interquartile range or extend from the lowest to the highest value if all values fall within the 1.5x interquartile range. Data are from 3 individual experiments with 3 replicates per condition, resulting in a total of 9 replicates per condition.

## Discussion

Bottom-up experimental approaches, in which a small number of bacterial species are mixed, have become popular for deriving principles of community dynamics and stability (6–12). Although co-culture experiments directly reveal winners and losers, they are often challenging to conduct. Specifically, co-culture experiments are often labor-intense and require phenotypic (morphological) or genotypic (fluorescence, antibiotic) markers to distinguish species, with the latter being often hard to introduce and limited in numbers (37–39). To overcome these limitations, we asked whether pairwise interaction outcomes and species rank order can be inferred from monoculture growth and supernatant assays without the need to mix species. While monoculture growth assays provide information on a species’ growth performance, supernatant assays provide information on secreted compounds that have inhibitory or stimulatory effects on opponents. Using a 4-species community, we found that monoculture growth is the most important factor in predicting competition outcomes, while supernatant effects allow fine-tuning of the species’ rank order. We further parameterised an agent-based model with our empirical data and conducted coculture experiments with 4-species and 3-species communities, to show that dynamics in multispecies communities match well the species rank order derived from monoculture and supernatant data with one important exception. The effect of inhibitory compounds is attenuated in multispecies communities, leading to an even higher predictive power of monoculture growth parameters.

For our community, we derived a set of three simple rules to predict species rank orders from monoculture growth and supernatant assays. It starts with the ranking of the species according to their monoculture growth performance (rule 1). Subsequently, species that show intermediate growth yet produce inhibitory compounds for other species are moved up in the ranking (rule 2), while inhibitory effects by slow-growing species are ignored (rule 3). The last rule applies because inhibitory compounds of slow-growing species do not reach a sufficiently high concentration to be effective in co-culture. Moreover, our results suggest that integrative growth parameters (1/T_mid_ and AUC) yield more robust predictions on competition outcomes than maximum growth rate. While maximum growth rate or yield are often used as fitness parameters (40,41), our findings are in line with other studies, showing that single growth parameters cannot adequately capture overall fitness across the duration of an experiment (13,38,42–44).

We do not claim that our rules are generally applicable to all bacterial communities, but we believe that our approach could be particularly useful for communities grown in batch culture under relatively homogenous conditions. Clearly, the applicability of our rules needs to be further tested using (i) media differing in their nutrient content, (ii) different environments including liquid and structured environments (16,34,45), and (iii) additional bacterial communities including both synthetic and natural bacterial assemblies. A key challenge of our approach is rule 2: how much should a species that suppresses the growth of others be moved up in the ranking? While this question was easy to address in our 4-species community with K consistently suppressing all others (in GIM), the situation will become complicated in larger communities with a diverse set of inhibitory interactions. In such situations, it might no longer be possible to precisely resolve rankings. However, rule 2 might still be useful to group species into categories of weak, intermediate, and strong competitors (10,31–33,38).

Our approach to predicting rank orders assumes that monoculture growth and supernatant assay data can be combined in an additive way. Whether pairwise interaction effects are additive or not will likely depend on the specific community and its members. Indeed, while some studies support additive effects (14,43,46), others show that non-additive effects in complex communities lead to new emergent properties (47–49). While we found most effects to be additive, our simulations identified ‘toxin absorption’ as a non-additive effect. Specifically, K is predicted to be the most competitive species in our community because it produces an inhibitory compound suppressing all other species. However, our simulations and 4-species community experiments showed that K is only the second-most competitive species after C. The reason for this mismatch is that the slower-growing species B and P show high per capita inhibitory compound uptake rates and thereby detoxified the environment for the fast-growing species C. This non-additive effect lowers the potency of inhibitory compounds in multispecies communities, and thereby affects the relative impact of our rules: they increase the weight of monoculture growth performance (rule 1) relative to supernatant effects (rule 2) in predicting competitive rank orders in multispecies community.

Another finding of our work is that competitive rank orders differ across growth media and the host. For example, P had a low competitive rank in GIM (3rd place), performed well in LB (2nd place), and was the top competitor in *G. mellonella* larvae (Figure 4). Several factors could contribute to such differences. First, each species has its nutritional preferences, which are better met in one of the two media. This could for example explain the markedly increased growth of C in GIM relative to LB. Second, the strength of negative interactions might be increased in medium that allows for higher growth because it favors higher production levels of inhibitory compounds. This could explain why K supernatant from GIM exerted higher negative effects on the other species than supernatant from LB. Third, niche diversity may differ across media, whereby lower niche diversity is predicted to intensify metabolic competition and niche exclusion. Finally, features of the host environment, such as spatial structure in tissue, reduced oxygen availability, and innate immunity can influence bacterial interactions. Altogether, our findings are in line with previous studies showing that a change in nutritional conditions can alter species interactions from positive to negative relationships (6,7,50) and suggest that special care must be taken when *in vitro* interaction data are used to forecast bacterial interactions in infections.

In conclusion, our work reveals that the combination of monoculture growth parameters, strong inhibitory effects from supernatant assays, and computer simulations can predict pairwise and multispecies interaction outcomes in a 4-species bacterial community. Next, it would be important to test whether our approach applies to more diverse bacterial communities and different environmental conditions. Moreover, identifying the actual compounds causing growth inhibition in the supernatant assays (e.g., toxins, siderophores, biosurfactants, amino acids, or molecules affecting environmental parameters such as pH) could further help to improve predictive power.

## Data availability

All raw data sets (https://figshare.com/articles/dataset/RawData_Experiments_Schmitz_etal/25187555) and simulation data (https://figshare.com/articles/dataset/RawData_Simulations_Schmitz_etal_/25187558) have been deposited in the Figshare repository.

## Supplementary information

Supplementary information is available online.

## Acknowledgments

We thank Richard Allen for his help with the statistics of this study. We also thank Alex Hall, Kayla King, Anna-Liisa Laine, and Roland Regoes for their scientific input.

## Compliance with ethical standards

Conflict of interest: The authors declare that they have no conflict of interest.

