## Supplementary Material (Figures and Table) for "Predicting bacterial interaction outcomes from monoculture growth and supernatant assays"

**Supplementary Information**
This file contains supplementary information for the article

### Predicting bacterial interaction outcomes from monoculture growth and supernatant assays

Désirée A. Schmitz[a], Tobias Wechsler[a], Ingrid Mignot[a] & Rolf Kümmerli [a]

[a] Department of Quantitative Biomedicine, University of Zurich, Zurich, Switzerland

The supplementary information comprises:

- 11 supplementary figures: Figure S1 – S11
- 1 supplementary table: Table S1

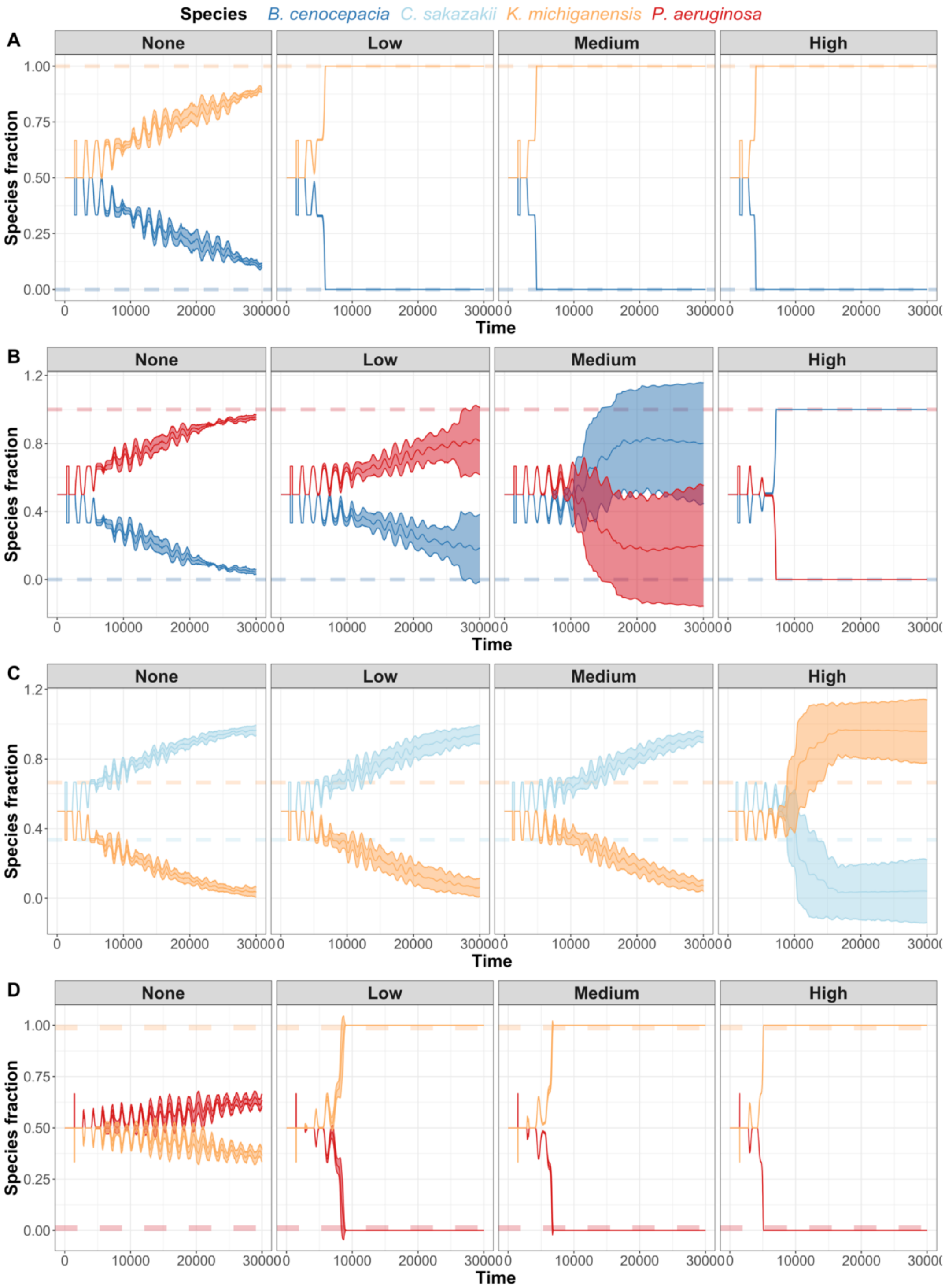


**Figure S1.** Pairwise competition outcomes from agent-based simulations in response to different toxin potencies. All simulations ran for 30,000 timesteps for the four pathogen interactions in which toxins were involved: (A) K versus B; (B) P versus B; (C) K versus C; (D) K versus P. The intrinsic growth parameter g was scaled relative to the mono growth data observed in GIM. Low, medium, and high toxin potency corresponds to a toxin threshold value of 2500, 2125, and 1750, respectively, which is the number of toxin molecules required to kill a cell. Hence, toxin potencies increase with lower threshold values. Lines show the mean and the standard deviation across 20 independent simulations. These simulations were carried out under conditions of high cell and molecule diffusion ($D$ = 5 μm$^2$ s$^{-1}$ and $\partial$ = 10 μm$^2$ s$^{-1}$ respectively) to match the experimental setup of a shaken culture. Dashed transparent lines show the results obtained in experimental co-cultures.

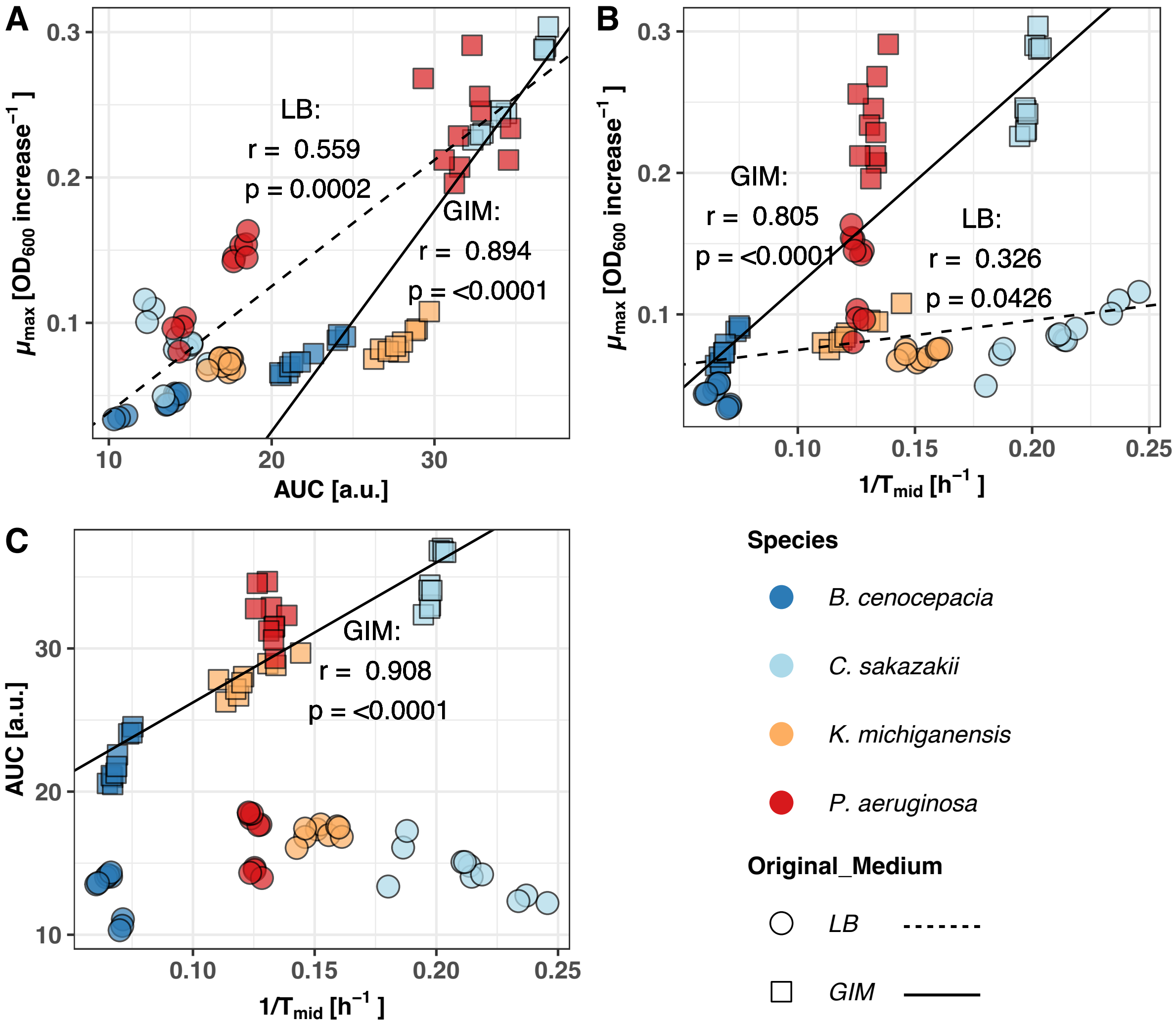


**Figure S2.** Correlation plots between the three growth parameters maximum growth rate ($\mu_{max}$), area under the curve (AUC, integral), and inverse of the time to mid-exponential phase ($1/T_{mid}$). Each color corresponds to a bacterial species, while the symbol of the data point corresponds to the type of medium (circles = LB; rectangles = GIM). The r-values indicate the Pearson correlation coefficient. The trendline is based on a linear regression analysis (dashed = LB; solid = GIM). Data are from 3 independent experiments, each featuring 3-4 replicates per condition, resulting in a total of 9-10 replicates per condition.

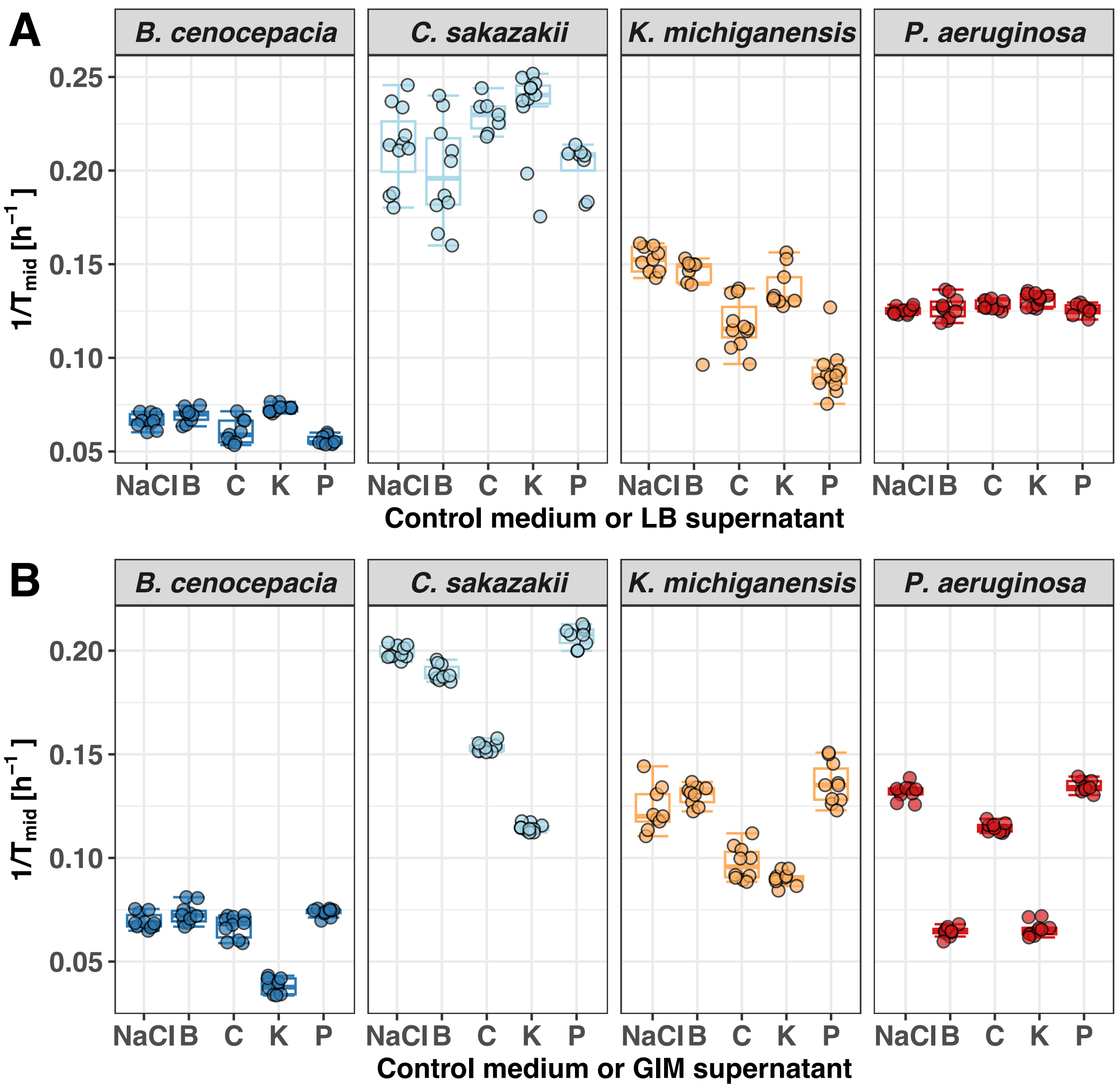


**Figure S3.** Boxplots show the absolute readouts of growth based on **$1/T_{mid}$** of each species in the conditioned medium (70% fresh medium + 30% spent supernatant) of the other species and a NaCl control treatment (70% fresh medium + 30% NaCl solution [0.8%]) in (A) LB and (B) GIM medium. Boxplots depict the median (line within the box) with the first and third quartiles. The whiskers cover 1.5x of the interquartile range or extend from the lowest to the highest value if all values fall within the 1.5x interquartile range. Data are from 3 independent experiments, each featuring 3-4 replicates per condition, resulting in a total of 9-10 replicates per condition.

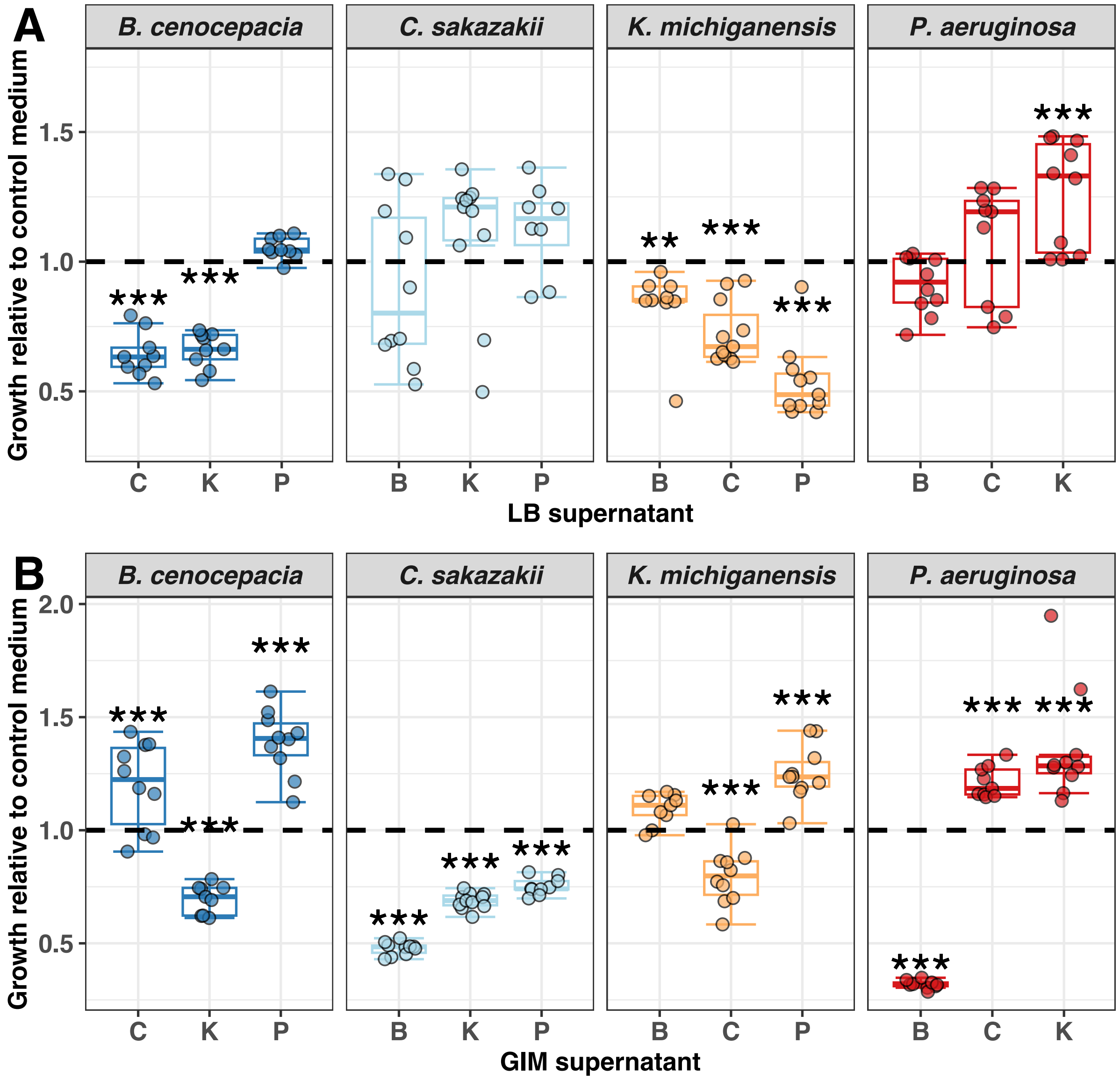


**Figure S4.** Boxplots show the relative growth based on $\mu_{max}$ of each species in the conditioned medium (70% fresh medium + 30% spent supernatant) of the other species both in (A) LB and (B) GIM medium, compared to a control treatment, depicted by the black dashed line (70% fresh medium + 30% NaCl solution [0.8%]). Relative growth was calculated by dividing the absolute $\mu_{max}$ (estimates from curve fits) in the supernatant treatments by $\mu_{max}$ in the control treatment (see Figure S5). Asterisks depict significant differences (alpha = 0.05) of a species' growth in the particular supernatant compared to its growth in the control medium using a linear mixed model with experimental block as random variable. Boxplots depict the median (line within the box) with the first and third quartiles. The whiskers cover 1.5x of the interquartile range or extend from the lowest to the highest value if all values fall within the 1.5x interquartile range. Data are from 3 independent experiments, each featuring 3-4 replicates per condition, resulting in a total of 9-10 replicates per condition.

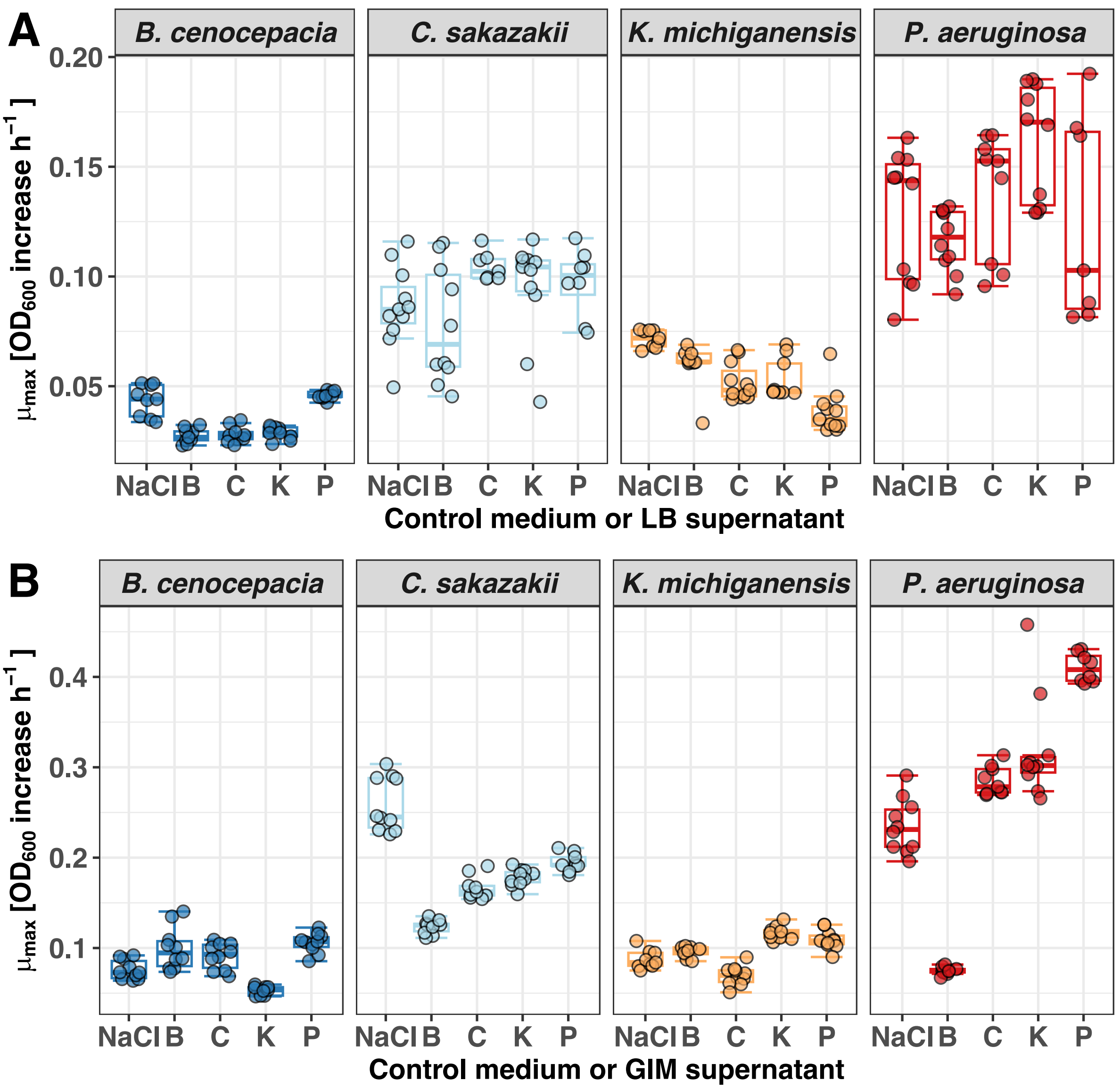


**Figure S5.** Boxplots show the absolute readouts of growth based on $\mu_{max}$ of each species in the conditioned medium (70% fresh medium + 30% spent supernatant) of the other species and a NaCl control treatment (70% fresh medium + 30% NaCl solution [0.8%]) in (A) LB and (B) GIM medium. Boxplots depict the median (line within the box) with the first and third quartiles. The whiskers cover 1.5x of the interquartile range or extend from the lowest to the highest value if all values fall within the 1.5x interquartile range. Data are from 3 independent experiments, each featuring 3-4 replicates per condition, resulting in a total of 9-10 replicates per condition.

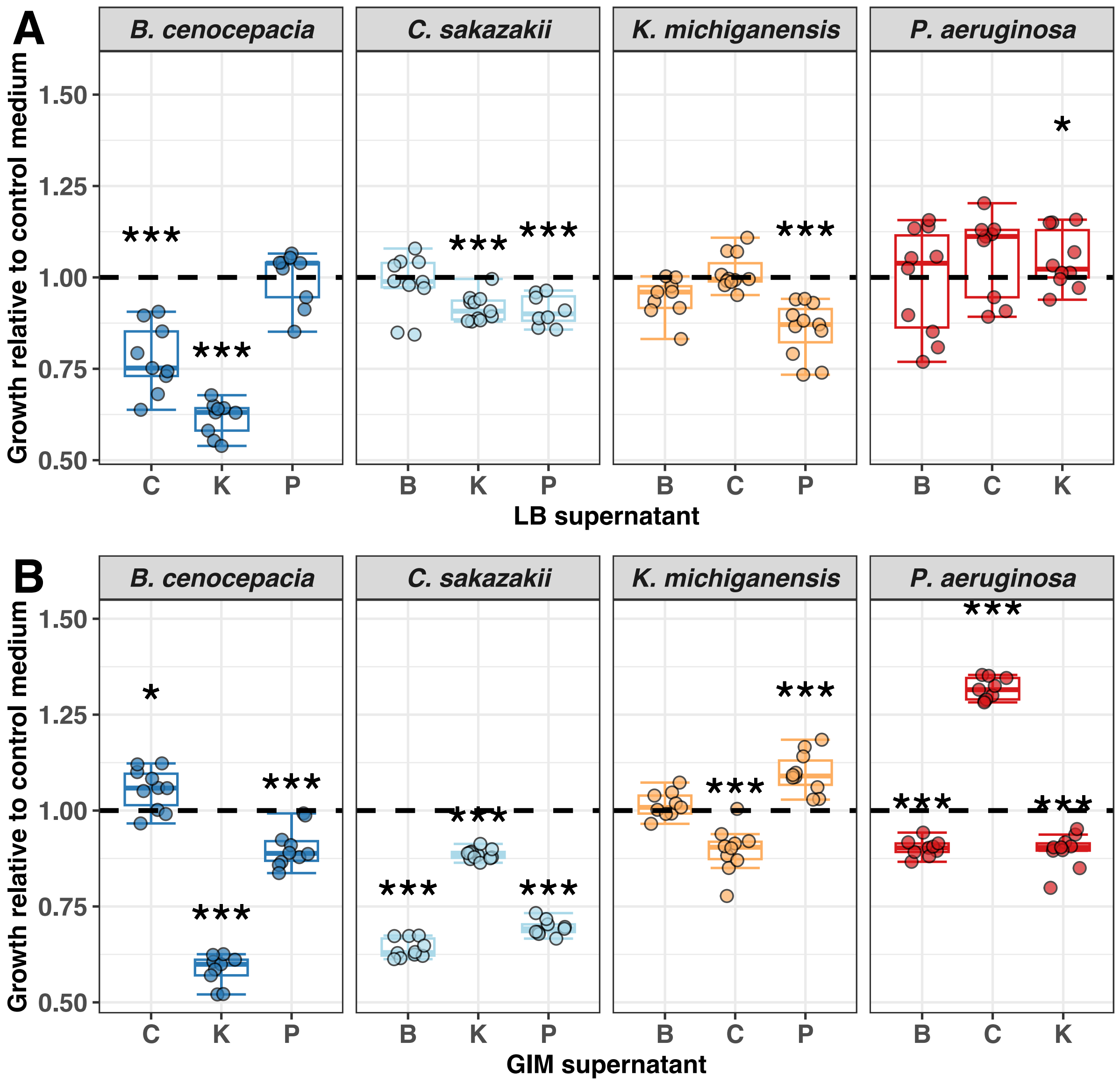


**Figure S6.** Boxplots show the relative growth based on the **AUC** (area under the growth curve) of each species in the conditioned medium (70% fresh medium + 30% spent supernatant) of the other species both in (A) LB and (B) GIM medium, compared to a control treatment, depicted by the black dashed line (70% fresh medium + 30% NaCl solution [0.8%]). Relative growth was calculated by dividing the absolute AUC (estimates from curve fits) in the supernatant treatments by the AUC in the control treatment (see Figure S7). Asterisks depict significant differences (alpha = 0.05) of a species' growth in the particular supernatant compared to its growth in the control medium using a linear mixed model with experimental block as random variable. Boxplots depict the median (line within the box) with the first and third quartiles. The whiskers cover 1.5x of the interquartile range or extend from the lowest to the highest value if all values fall within the 1.5x interquartile range. Data are from 3 independent experiments, each featuring 3-4 replicates per condition, resulting in a total of 9-10 replicates per condition.

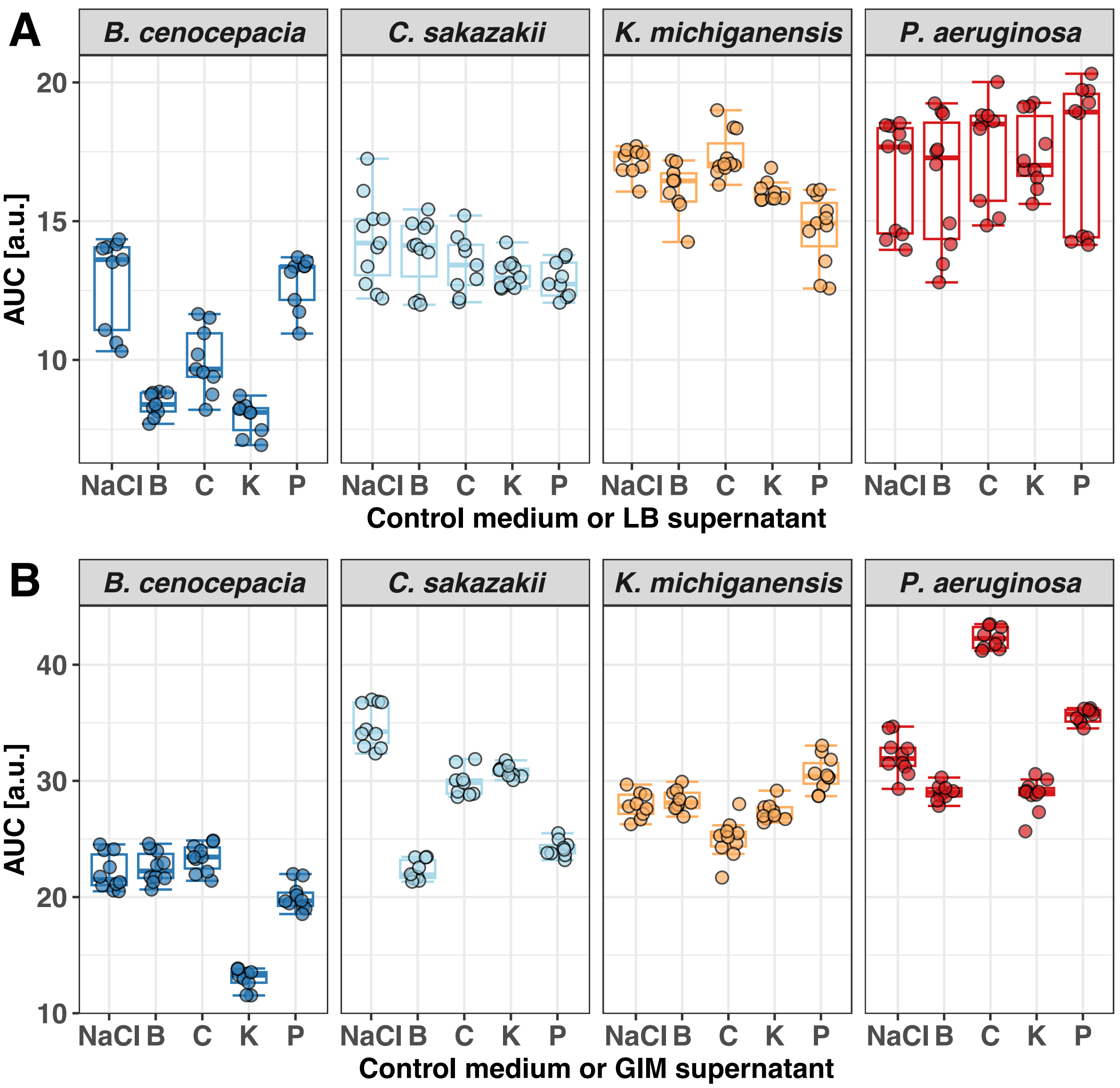


**Figure S7.** Boxplots show the absolute readouts of growth based on **AUC** (area under the growth curve) of each species in the conditioned medium (70% fresh medium + 30% spent supernatant) of the other species and a NaCl control treatment (70% fresh medium + 30% NaCl solution [0.8%]) in (A) LB and (B) GIM medium. Boxplots depict the median (line within the box) with the first and third quartiles. The whiskers cover 1.5x of the interquartile range or extend from the lowest to the highest value if all values fall within the 1.5x interquartile range. Data are from 3 independent experiments, each featuring 3-4 replicates per condition, resulting in a total of 9-10 replicates per condition.

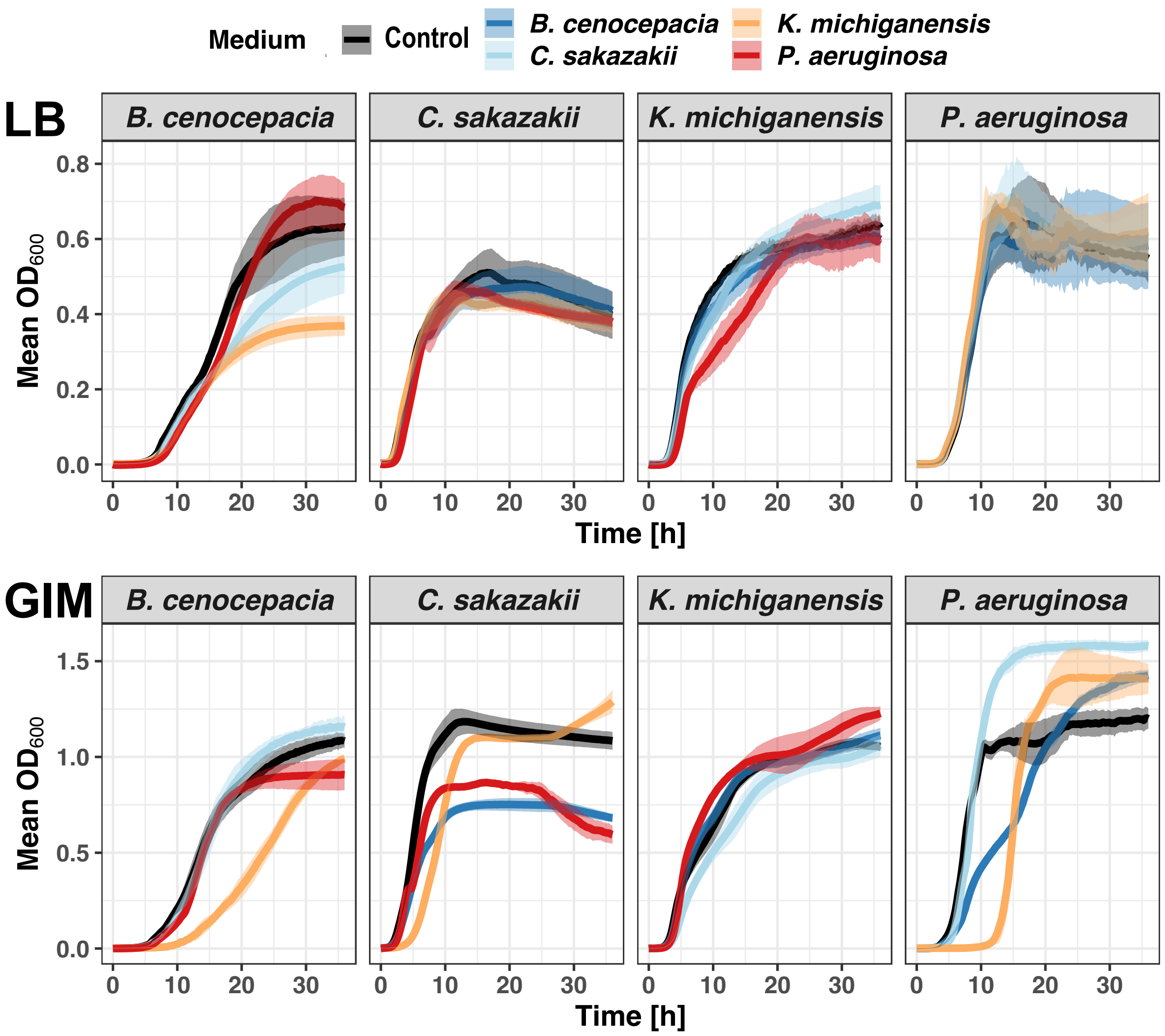


**Figure S8.** Growth curves show the mean $OD_{600}$ measurements in conditioned medium (70% fresh medium + 30% spent supernatant) of the other species and a NaCl control treatment (70% fresh medium + 30% NaCl solution [0.8%]) in (A) LB and (B) GIM medium. Each colored curve stands for the conditioned medium of a specific species (see legend above panels) and the black line is the control. Shaded areas depict the standard deviation. Data are from 3 independent experiments, each featuring 3-4 replicates per condition, resulting in a total of 9-10 replicates per condition.

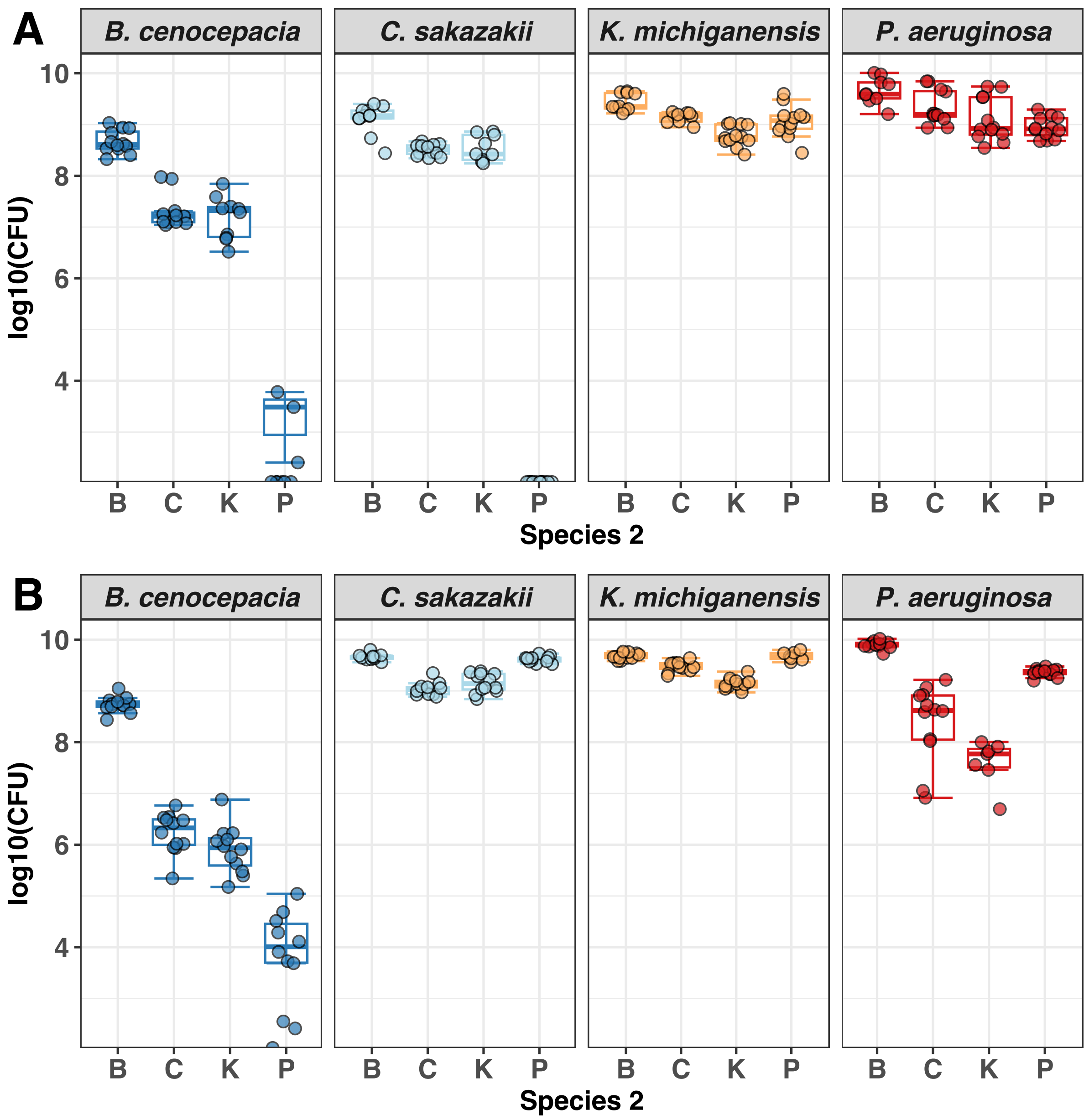


**Figure S9.** Boxplots depict the CFU of the focal species (indicated in the header of each panel) in monoculture and in co-culture with each of the other species after 24 h in (A) LB and (B) GIM medium. The CFU-values of monocultures are corrected (divided by two) to account for the fact that the focal species inoculum was double in mono- compared to co-cultures. Boxplots show the median (line within the box) with the first and third quartiles. The whiskers cover 1.5x of the interquartile range or extend from the lowest to the highest value if all values fall within the 1.5x interquartile range. Data are from 6 individual experiments with 2 replicates per condition, resulting in a total of 7-12 replicates per condition (note in a few cases sample size was <12 because obtaining countable colonies for both species in co-culture was very difficult).

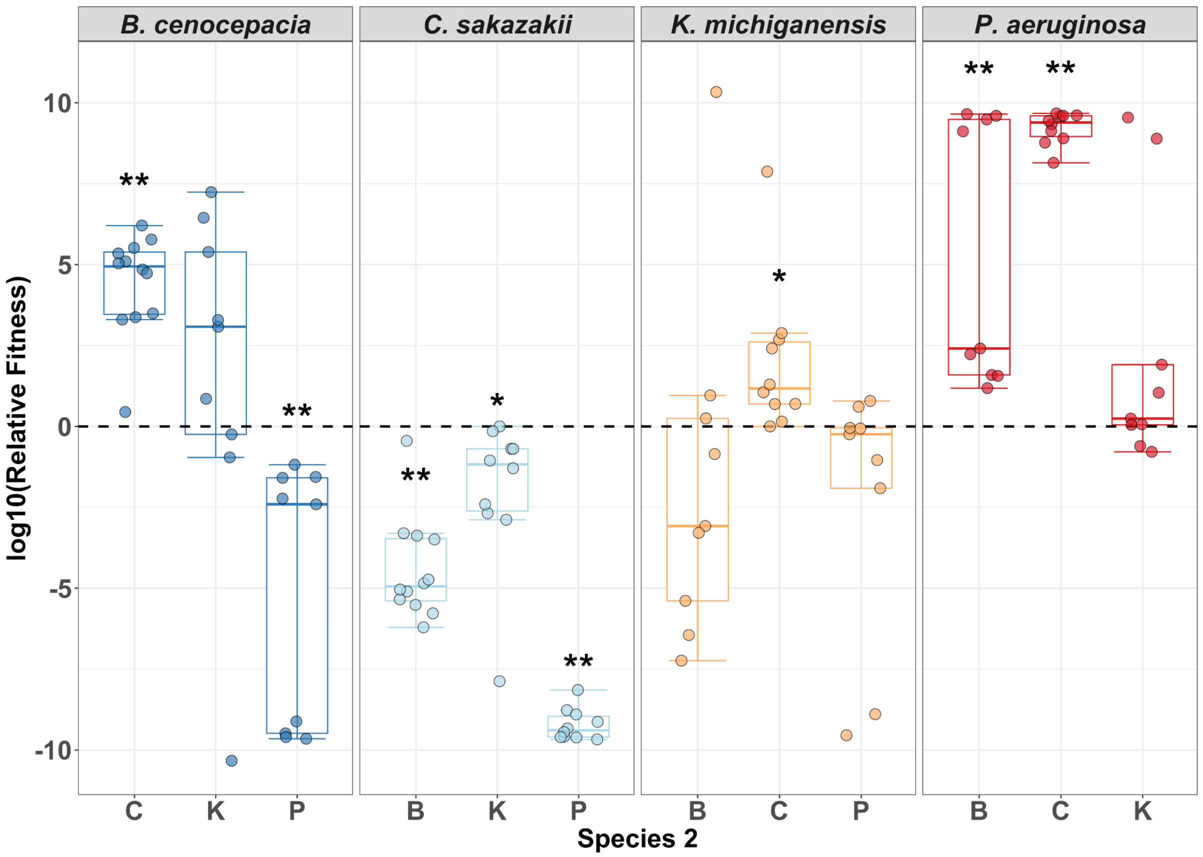


**Figure S10.** Relative fitness in pairwise competitions 12 hours post-infection in the larvae of *Galleria mellonella* from Schmitz et al. (2023)*. Relative fitness was calculated using Wrightian fitness with the number of cells injected into a larva at the start and the number of colony-forming units counted 12 hours post-infection (see methods in Schmitz et al. 2023 for details). The dashed black line indicates equal fitness of both species in co-infections, while negative and positive values represent decreased and increased fitness of the focal pathogen, respectively, relative to the co-infecting pathogen. Boxplots depict the first and third quartiles and the median line in between, while its whiskers cover 1.5x of the interquartile range or extend from the lowest to the highest value if all values fall within the 1.5x interquartile range. Each dot represents an individual larva. Asterisks depict significant differences (alpha = 0.05) of a species' fitness relative to a competitor against the null hypothesis that relative fitness is the same (black dashed line) using one-sample Wilcoxon rank tests. Data are from 4-5 individual experiments, each featuring 2-3 larvae per treatment, resulting in a total of 8-12 larvae per treatment. In the main text, we compare the competitive ranking of the four pathogens *in vitro* and *in vivo* (based on the data depicted in this figure) and only find a moderate match. Importantly, we used the same methods for the *in vivo* and *in vitro* assays by mixing equal amounts of two pathogens and enumerating their CFU after a defined time point. While the basis for the fitness measurement is thus the same, fitness and rank orders are different *in vitro* versus *in vivo*. Several factors can contribute to these differences. For example, the host environment offers more spatial structure, different biotic and abiotic factors as well as dynamic host factors, which are absent in the shaken, liquid cultures of the *in vitro* experiments. Our results thus suggest that special care must be taken when *in vitro* interaction data are used to forecast bacterial species interactions in infections.

* Schmitz DA, Allen RC, Kümmerli R. Negative interactions and virulence differences drive the dynamics in multispecies bacterial infections. Proceedings of the Royal Society B 2023. 290: 20231119.

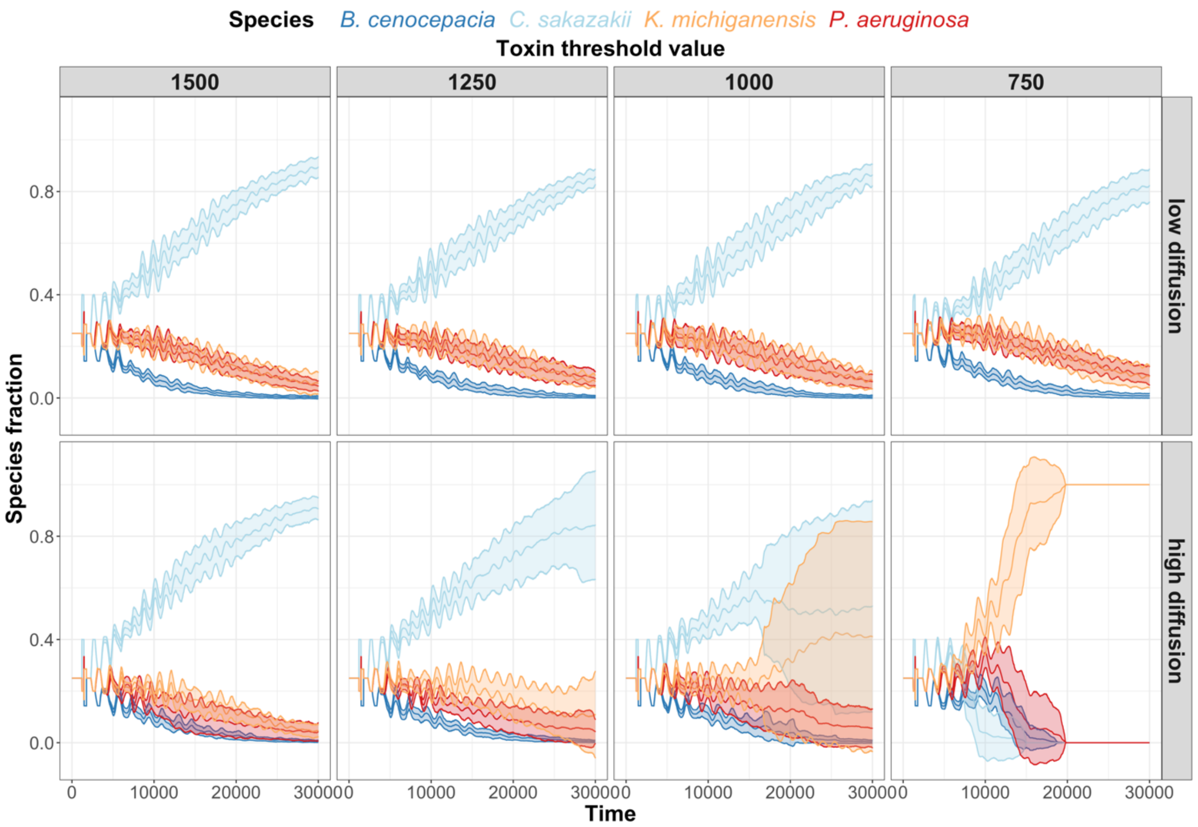


**Figure S11.** Variation in 4-species community dynamics in response to K toxin potency and diffusion. All simulations ran for 30,000 time steps with the four pathogens starting at equal fractions. Toxin potency is defined by the toxin threshold value, which is the number of toxin molecules required to kill a cell. Hence, toxin potencies increase with lower threshold values. Low diffusion (cell diffusion 0.0 μm$^2$ s$^{-1}$, toxin diffusion $\partial$ = 0.1 μm$^2$ s$^{-1}$) corresponds to a structured environment. High diffusion (cell diffusion 5.0 μm$^2$ s$^{-1}$, toxin diffusion $\partial$ = 10.0 μm$^2$ s$^{-1}$) corresponds to an unstructured environment as implemented in our experiments with shaken cultures. Lines show the mean and the standard deviation across 20 independent simulations.

**Table S1.** Values of pH for 30% supernatant + 70% original medium for all four species as well as in both media. SN = supernatant.

| Medium | Species | pH of pure SN | pH of 30% SN + 70% original medium |
|---|---|---|---|
| **LB** (pH 6.95) | B | 8.48 | 7.54 |
| | C | 8.31 | 7.4 |
| | K | 8.95 | 7.87 |
| | P | 8.05 | 7.13 |
| **GIM** (pH 6.01) | B | 7.40 | 6.40 |
| | C | 5.32 | 5.76 |
| | K | 5.22 | 5.60 |
| | P | 7.35 | 6.22 |
